# Cultivated plants in the demographic projections of the global carbon budget

**DOI:** 10.64898/2026.01.14.699272

**Authors:** Arnaud Muller-Feuga

## Abstract

Global carbon budgets attribute an incomplete role to cultivated plants due to the exclusion of annual crops, which are not considered to result in net carbon accumulation. Using FAO agricultural and forestry production statistics, this study re-evaluates the amounts of CO₂ captured, stored, and restituted by cultivated plants, explicitly including annual crops, forage crops, forest products, and non-marketed plant parts.

A stoichiometric and probabilistic approach is developed to describe the temporal dynamics of photosynthetic carbon capture and restitution. The results show that the net balance of cultivated plants has constituted a sink of 3.0 ± 0.7 GtCO₂/year on average since 1970, with an average half-life of 10.7 ± 2.9 years. This annual contribution of cultivated plants to atmospheric carbon storage in organic form is sufficient in both quantity and duration to be included in carbon budgets. Among them, annual crops accounted for approximately 27% of total carbon sequestration with retention periods of 6 years, thus invalidating their exclusion from the carbon budgets.

The strong correlation observed between the global human population size and the elements of the carbon budget allows for projections up to the end of the 21st century. These projections, adjusted by the United Nations’ median population scenario, suggest that the carbon sink of cultivated plants would reach zero around the time of the global population peak before reversing. The carbon stocks built by these plants increased by 43 GtC between 1970 and 2024 and would increase by a further 21 GtC by the time of the population peak before declining. Around the same time, the carbon sink constituted by the ocean and unexploited land areas would peak at 25 GtCO_2_/year, and that constituted by the atmosphere at 24 GtCO_2_/year, with atmospheric CO_2_ concentrations reaching 600 ppm. Emissions from fossil fuel combustion would peak at 48 GtCO_2_/year by this same timeframe before declining.

These results highlight the significant role of cultivated plants in the carbon cycle and show that excluding annual crops leads to a substantial underestimation of continental carbon sinks. They call for a revision of carbon accounting methods and a full integration of agriculture and livestock farming into global carbon budgets.

**Key points:**

- Although small compared to other carbon sinks, cultivated plants must be included in carbon budgets because they build up a significant carbon stock.
- When demography is used to modulate projections, most elements of the carbon budget reach their maximum at the same time as population, while the cultivated plants carbon sink declines and turns a source.

## 1. Introduction

Because greenhouse gases in the atmosphere could influence Earth’s climate, attention has been focused for several decades on exchanges involving CO_2_. The global warming that has been occurring for half a century at a rate of 0.19°C every 10 years (Met Office Hadley Centre, 2022) is thought to be a consequence of the release into the atmosphere of CO_2_ produced by the combustion of fossil fuels since the beginning of the industrial era. The atmospheric concentration of CO_2_ (ACC) increased from 270 parts per million by volume (ppm) in the mid-19th century to 420 ppm in 2024. According to IPCC (2021), CO_2_ emissions from human activities during the decade 2010-2019 were due to the combustion of fossil fuels (81-91%), with the remainder being the net CO_2_ flux related to land-use change and land management. These emissions are estimated to have three destinations: 46% accumulated in the atmosphere, 23% was absorbed by the ocean, and 31% by terrestrial vegetation.

Plants are cultivated to meet humanity’s needs for food (cereals, vegetables, fruits, etc.), textiles (cotton, linen, hemp, etc.), ornamental purposes (tobacco, grapes, flowers, etc.), heating, and construction. They achieve the direct capture of atmospheric CO_2_, then the polymerization of carbon into biomass according to the process of photosynthesis at the origin of life on Earth. Carbon capture and storage in marketed plant products from agriculture, livestock farming, and forestry was estimated (Muller-Feuga, 2024a) at 21 billion tonnes of carbon dioxide per year (GtCO_2_/yr) in 2022. Subsequently, the non-commercial parts of these products were considered (Muller-Feuga, 2024b), which attributed 41.0 ± 0.6 GtCO_2_/yr to whole cultivated plants, with a dry weight-weighted average storage duration of 26.3 ± 2.0 years in 2022. The analysis was complemented by an examination of the evolution of carbon capture and restitution over the past 50 years (Muller-Feuga, 2025a), which showed that plant cultivation constitutes an average carbon sink of 39.9 GtCO_2_/yr removed from the atmosphere during the ten years preceding 2022. These overestimated results are corrected below.

The differences between these figures and those in the literature appear to be due to the failure to consider annual crops in carbon budgets. We noted a recommendation to this effect among the methods and guidelines for researchers (IPCC, 2006; 2019). The increase in biomass of annual crops would be equal to the biomass losses of the same year, and there would therefore be no net carbon accumulation, as only perennial woody crops should be considered. This has the effect of significantly reducing the proportion of plants included in carbon budgets.

In reality, the carbon in cultivated plants is not released quickly except in the extreme case where the crop is burned. Even in this situation, the underground parts, which represent a substantial portion of the biomass, persist for several years, even decades. The death of the harvested annual plant does not mean the disappearance of the carbon it has accumulated. Grass products, representing two-thirds of global agricultural production, have indefinite storage life under the right humidity and temperature conditions, either as grains or after processing into dry products. Long storage periods before marketing allow for price stabilization, long-distance transport, and the creation of strategic reserves by populations facing crises or living far from production areas.

It is important to distinguish between carbon stocks and the carbon capture and restitution flux of atmospheric CO_2_, the latter being the time derivative of the former. Carbon stocks, measured in tonnes (tC), are the quantity accumulated at a given location and time. Only the flux, expressed in tonnes of carbon dioxide (tCO_2_/year), should be considered in annual carbon budgets. By convention, “sink” fluxes, which remove carbon from the atmosphere, are negative, and “source” fluxes, which release carbon into the atmosphere, are positive. The plant sink is proportional to the amount of plant biomass cultivated and then harvested annually for later use elsewhere. The plant source is the return to the atmosphere of this biomass after mineralization.

Thanks to their intrinsic capacities and conservation techniques, annual plant biomass persists well beyond the harvest year. Here we provide the necessary updates and corrections to determine the extent to which the exclusion of these factors impacts the carbon footprint. We also offer a projection up to the end of the century that incorporates these corrections.

## 2. Materials and Methods

The synthesis of plant organic matter involves a chain of enzymatic reactions, notably involving carbonic anhydrase and rubisco, for the entry of carbon dioxide (CO_2_) from the atmosphere in dissolved form into the plant cell, followed by a series of reactions modulated by visible light. The biomass thus formed serves as a substrate for heterotrophic organisms, which return carbon to the atmosphere as CO_2_ through respiration or fermentation. These polymerization and mineralization reactions are grouped under the same reversible chemical equation (1)

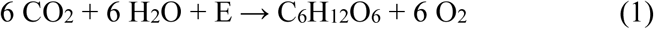

where E is the visible light energy in the direction of photosynthesis (from left to right) and the metabolic or combustion energy in the direction of respiration (from right to left).

Equation (1) expresses that CO_2_ consumption, oxygen production, and organic matter production in the form of hexoses correspond molecule for molecule. Hexose is the basic building block of plant matter, and its quantity can be measured by the dry matter (dm) of plant biomass according to the stoichiometry of the chemical reaction. The carbon-to-dry matter ratio (C/dm) varies with the type of biomass and should be adjusted for each case.

Although the energy conversion efficiency of solar radiation into biomass is on the order of a few percent, the resulting plant production is an important carbon sink. Controlling all or part of the food chain, agriculture, livestock farming, forestry, hunting, fishing, and aquaculture feed, clothe, warm, shelter, and entertain humanity, among other things. Here, we consider the net primary production after autotrophic respiration of these activities, which capture carbon from the atmosphere and store it as biomass. As such, they are involved in the carbon cycle and its budget.

### 1. The quantities of carbon captured and stored

The calculation of the quantities of carbon mobilized by crops is based on the FAO’s 2024 statistics (n.d.), which describe the products marketed from agriculture, livestock farming, and forestry. The amount A_n_ of CO_2_ absorbed by a set b of plant biomass with fresh weight P_n_ is equal to the sum of the dry weights of the crops multiplied by the amount of CO_2_ per dry weight (CO_2_/dm), denoted k, according to formula (2), where WC_n_ is the water content of the biomass P_n_ documented by various cross-referenced sources.

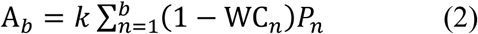

The carbon content C/dm varies with the species and, within the species, with the organ. The biomass molecules have a C/dm of 40% for glucose, 44% for cellulose, 64% for hemicellulose, and 66% for lignin. According to Ma S. et al. (2017), the ratios (C/dm) of reproductive organs, roots, leaves, and stems are 45.0%, 45.6%, 46.9%, and 47.9%, respectively. The greatest difference in carbon content is between annual and perennial plants. Regarding reproductive organs, which are the main crop products, those of annual plants contain 42.5% carbon and those of perennial plants 48.6%. We will use these values for crops and forage. Regarding forest products, which consist primarily of woody stems, the main distinction is between conifers (50.5%) and non-conifers (48.2%).

### 2. Carbon capture and storage durations

The carbon history of agricultural and forestry products is divided into the period of carbon capture by photosynthesis (CP) and the period of carbon restitution by mineralization (RM). The harvest marks the transition between these two periods of anabolism (CP) and catabolism (RM), which involve the same quantity of carbon. During the first period (CP), which separates the beginning of plant growth, whether through sowing, planting, or the previous harvest (n-1), from the harvest (n), the plant carbon pool is built up through photosynthetic capture of CO_2_ from the atmosphere. This anabolic period lasts from a few months for annual plants to several decades for trees. The second period of catabolism (RM) begins immediately after harvest, during which the captured carbon is restituted to the atmosphere as CO_2_ through respiration, combustion, or fermentation. During this period, marketed biomass is stored on shelves, in silos, and in warehouses, transported, and processed before being used by the end consumer.

The quantities of CO_2_ captured and then restituted are calculated based on the carbon per dry matter ratio (C/dm) of each biomass. The CP and RM durations are the weighted averages for each of the three biomass groups: crops, forage, and forestry. The dispersion of the results is expressed by their standard deviation (SD). The maximum durations of the CP and RM periods are determined for each biomass based on the most relevant data from the literature. From 1960 to 2020, only harvests from years that are multiples of 10 are considered, using the CP and RM values obtained for 2024.

In addition to temperature, the RM shelf life of food depends on its water content (WC), which can be reduced by various techniques such as drying, brining, and vacuum packing. We thus distinguish (Muller-Feuga, 2025b) between fresh fruits and vegetables, for which the WC is between 75 and 90% and for which the RM is a few days to a few weeks; semi-preserved products, for which WC is between 30 and 75% and the RM is a few weeks to a few months; and dry products such as cereals, pasta, and biscuits, for which the TE is between 0 and 30% and for which the RM is a few months to indefinitely.

Since the nutritional value and food safety (freshness) decrease with age, consumers are guided in their choices to ensure food safety and limit waste. Recommended storage times (Use By, Best Before, etc.) are mandatory information displayed on food packaging. As this information is not always available, we considered the maximum recovery time to be a function of the water content WC according to regression (3) for which R² = 0.4.

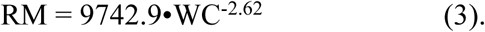

Freshly harvested fruits and vegetables have a shorter transport and marketing time to preserve their taste and freshness. Temperature control during transport and storage significantly increases the time between harvest and consumption. This shelf life varies from a few days to a few weeks at room temperature or in a refrigerator (-5°C). It can last up to 3 months, allowing them to last through the winter. In a freezer (-18°C), food generally keeps for a year or more.

### 3. Non-marketed parts

It is also necessary to consider non-marketed but inherent parts of crops, such as leaves and stems for the aboveground portion, and roots and exudates for the underground portion. The half-lives of necromass, consisting of aboveground and underground parts remaining in place after harvest, correspond to the average residence time of carbon in the soil before mineralization by decomposer organisms. For forests, these retention times vary between 0.9 and 152 years (Wang J. et al., 2017). They increase with latitude (x4) and decrease with temperature (x10) and precipitation (x4). We have used temperate regions as an average, for which the maximum residence time of organic carbon is 40 years for soils of crops and grasslands, and 75 years for soils of harvested forests (Balesdent and Recous, 1997; Pellerin et al., 2019).

The weights of the aboveground parts of crops are calculated based on an average harvest index (aboveground/commercial part) of 0.42 (Hay R., 1995). The root system is estimated to represent 30% of the biomass of cultivated plants, 65% of grassland plants, and 15% of the total biomass of trees (Blume et al., 2015). Consequently, the ratios of whole plant weight to commercial part are 1.72, 2.07, and 1.57 for crops, forage, and forest products, respectively. The CO_2_/dm ratio of commercial parts is extended to whole plants.

The quantities of atmospheric CO_2_ mobilized by cultivated plants in agriculture, livestock farming, and forestry are updated here in terms of both the amount captured and the duration of carbon sequestration. For this purpose, the productions are converted into anhydrous products according to formula (2), then multiplied by the ratio of whole plants / commercial parts, then multiplied by their CO_2_/dm ratio.

### 4. Probabilistic approach to the plant carbon cycle

The photosynthetic growth of plants bearing logs, fruits, and vegetables during the carbon capture period (CP) is measured by the weight gain per unit time, denoted dP/dt. This is a real random variable whose distribution generally exhibits an increasing sigmoidal shape (e.g., Vanclay, 1994; Karadavut et al., 2008; Tijero et al., 2021). In what follows, we consider this distribution function to be the integral of a increasing normal probability density function.

After harvest, the polymerized carbon is mineralized and returned to the atmosphere as CO_2_ through food digestion, fermentation, or combustion. This restitution period (RM) varies from a few days for perishable fruits and vegetables to several centuries for timber and soils. Organic matter, whether stored on shelves, in bulk, or in the soil, is ingested, fermented, or burned, resulting in weight loss. The distribution of this weight over time follows a decreasing sigmoidal curve, which can be approximated by the integral of a negative normal density law.

For both carbon capture and restitution, the normal density law is chosen here for its continuity, ease of manipulation, and its ability to accurately describe phenomena influenced by numerous factors, none of which is dominant. The probabilistic simulation of the temporal distribution of atmospheric CO_2_ mobilized in plant matter, described in Muller-Feuga (2024b, 2025a), is reproduced here, updated, and corrected. We assume that the timing of atmospheric CO_2_ capture and restitution is represented by the following expressions, for capture:

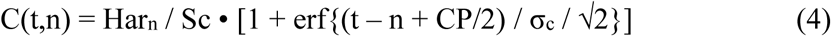

and for restitution:

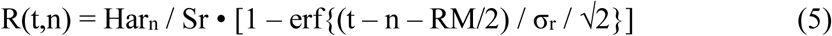

where t is the time in years, n is the harvest year, Sc and Sr are adjusted so that the sums of the column vectors equal the harvest of year n, denoted Har_n_, erf is the error function of the normal distribution, and σ_c_ and σ_r_ are the camber or standard deviations. If t ≤ n, the uptake duration CP applies, and the erf function is added (4). If t > n, the restitution duration RM applies, and the erf function is subtracted (5).

The reduced distribution of CO_2_ mobilized by the 2024 global harvest (Figure 1) results from the integration of probability densities. A significant difference between the CP and RM durations was observed (Muller-Feuga, 2024b, 2025b), with the former being shorter than the latter. Consequently, the distribution exhibits a strong asymmetry. Since normal distributions are centered, the mean and median carbon storage half-lives (CSD) are equal to the sum of half the maximum capture duration (CP/2) and half the maximum restitution duration (RM/2) according to formula (6).

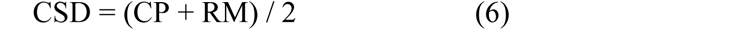

**Figure 1:**
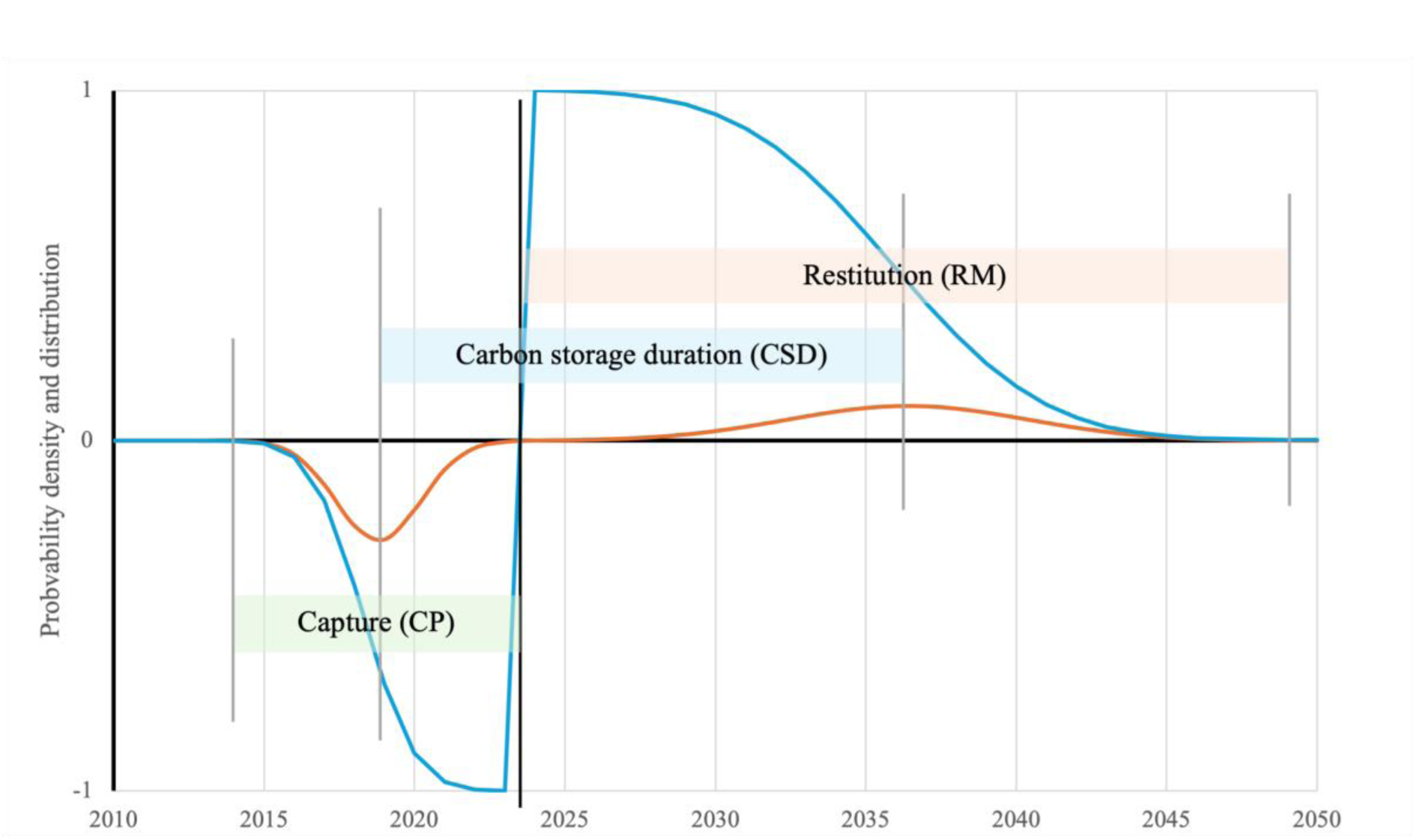
Reduced chronologies of the probability densities (red) and cumulative distribution (blue) of the carbon captured by photosynthesis and then restituted by mineralization, for the 2024 global harvest. Capture is negative and restitution is positive. The periods CP, RM, and DSC are illustrated.

This probabilistic approach is applied to cultivated plants with a single harvest, such as annuals and trees felled for firewood and construction timber, and to perennial plants, such as sugarcane, tea, coffee, cocoa, oil palm, rubber trees, fresh and dried fruits, etc., harvested through multiple annual crops. In the latter case, the capture period CP covers the plant’s growth, which varies between 2 and 80 years depending on the species.

The theoretical chronology of carbon capture and storage by annual plants, simulated by equations (4) and (5), allows us to construct three 173x173 square matrices, which we named [C(t,n)] for capture, [R(t,n)] for restitution, and [C(t,n) – R(t,n)] for the complete capture-restitution cycle, where t and n are between the years 1960 and 2133. The row vectors t contain the quantities of CO_2_ mobilized by successive harvests, and the column vectors n contain the quantities captured and then restituted from the harvest Har_n_ as a function of time t. Only the diagonal area for increasing t and n of these three matrices is non-zero.

The global capture in year n, denoted AW_n_, is equal to the sum of the row vector t of the matrix [C(t,n)], which is written (7).

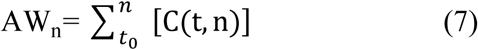

Similarly, the global restitution of year n, denoted EW_n_, is equal to the sum of the row vector t of the matrix [R(t,n)], which is written (8).

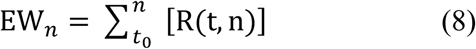

The net capture minus restitution balance of year n is equal to the sum of the row vector t of the matrix [C(t, n) – R(t,n)], which is written (9).

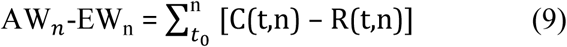

The year-by-year accumulation of these net balances, positive or negative, constitutes the carbon stocks built up by cultivated plants from t0 to n, denoted CS, which have the following matrix expression (10).

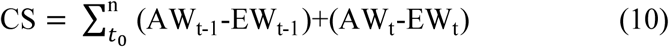

### 5. The carbon budget

Global CO_2_ absorption by cultivated plants photosynthesis (denoted AW) and global CO_2_ emissions from the digestion, fermentation, or combustion of these plants (denoted EW) constitute the anthropogenic components of global atmospheric carbon budgets. Atmospheric CO_2_ mass variations (denoted ACV) and emissions from fossil fuel combustion (EFOS) complete the carbon exchanges between the continents, the atmosphere, and the oceans. The unknown, denoted OS+, in equation (11) includes the contribution of the oceans as well as that of uncultivated land. The balance of these exchanges describes the equilibrium between these five components, where sources are positive and sinks are negative.

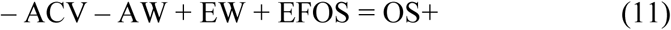

The estimation of AW is based on the stoichiometry of reaction (1) in the left-to-right direction of photosynthesis, while that of EW results from the same reaction in the right-to-left direction of mineralization. This approach does not rely on numerical models based on satellite or field-measured data, but only on agricultural and forestry production data reported by governments and compiled by the Food and Agriculture Organization of the United Nations (FAO).

### 6. Projections of carbon budget elements

There is a proportional relationship between the size of the world’s population and the consumption of fossil fuels EFOS, on the one hand, and plant-based products AW, on the other, since both are intended to meet humanity’s needs. We assume that the increase in atmospheric CO_2_ mass ACV also depends on the world population, since it is directly influenced by anthropogenic emissions EFOS and EW. This assumption is used to anticipate the evolution of carbon budgets until the end of the century.

The UN median scenario from July 2024 (United Nations, 2024) is chosen as the modulator due to its higher and more recent probability. Following industrialized countries, the transition to two children per couple, stabilizing the population, should spread to all continents in the coming decades. The population would exceed 10 billion inhabitants in 2060 and peak in 2084 before declining. The OS+ contribution, which includes that of the ocean and undeveloped land areas, results from equation (11).

## 3. Results

### 1. Commercial crop products 2024

The 160 global crop products are listed by the FAO (n.d.) under the heading “Products,” then “Crops and livestock products” in the agricultural production statistics. They include herbaceous and shrubby plants cultivated for food (cereals, vegetables, fruits, etc.), textiles (cotton, flax, hemp, etc.), or ornamental purposes (tobacco, grapes, flowers, etc.). These products are listed in descending order of fresh weight in Annex 1 at the end.

Whole plants of commercial crop products had sequestered 14.5 ± 0.3 GtCO_2_. The largest contribution was from dried and preserved products, accounting for 87% of this sequestration. Annual plants contributed 75 % to atmospheric carbon sequestration by crops, of which 61% was due to cereals. The average carbon-weighted CP, RM, and CSD of crops were 4.9 ± 0.21, 20.6 ± 1.0, and 12.7 ± 0.6 years, respectively. Excluding annual plants by considering only perennials means neglecting a major component of carbon budgets.

### 2. Forage 2024

Forage consists mostly of annual plants, the harvested crops of which are stored for 0.8 to 3 years to allow them to mature and pass through unproductive periods. They are transformed into meat, milk, offal, eggs, honey, etc., by the animals that consume them and use their carbon for their structure and metabolism. Carbon is restituted as CO_2_ through livestock respiration and the mineralization of their excrement and non-food products (hides, wax, silk, etc.). It lasts between 40 days (eggs) and 100 years (beeswax).

The 48 global livestock products are listed as quantities of meat, milk, and eggs consumable by humans, expressed in tonnes. The statistics described in the “Production – Quantity” and “ Livestock Primary “ sections are included in Annex 2 and are ranked in descending order of dry weight of forage. These products are primarily intended to supplement the human diet with high-quality protein and a range of micronutrients such as vitamins A, B-12, riboflavin, and minerals. They result from the conversion of plants from grasslands, foliage, and crops by ruminant and monogastric animals. Protein levels vary between 3.2% for milk and 32% for meat products. Feed conversion ratios (dry weight of feed/fresh weight gain of livestock product) vary between 2 for poultry and fish fed dry feed and 6 for cattle fed aboveground plants.

According to Mottet et al. (2017), global livestock consumed 6 Gt/year of dry matter forage in 2010. This yields an average conversion ratio of 4.34 across all species. Assuming that it is stable from year to year, we applied it to the 2024 livestock to calculate anhydrous forage, then the CO_2_/dm ratio, then the whole plant/commercial part of annual plants ratio to obtain the CO_2_ mobilized.

A portion of the crops described in the previous section is intended for animal feed. This mainly consists of cereals (wheat, maize, sorghum, oats, etc.) used in livestock feed. To avoid double counting, a 14% reduction is applied to forage calculated based on marketed animal products. This represents the portion of forage consumable by humans, according to the authors cited above. Thus, the assimilable parts of forage plants for global livestock would have sequestered 8.7 GtCO_2_/year of carbon in 2024, or 18.2 ± 1.6 GtCO_2_/year for whole plants. The carbon-weighted average CP, RM, and DSC of forage plants were 1.18 ± 0.08, 0.68 ± 0.04, and 0.93 ± 0.06 years, respectively.

### 3. Forest products 2024

Statistics on global timber production are available in the “Forestry” section of FAO’s “Data” (n.d.). These “Forestry production and trade” figures for fuelwood, sawlogs, pulpwood, and industrial wood, expressed in m³, are converted to dry tonnes and listed in descending order in Annex 3. A density of 0.6 t/m³ is used for conifers and 0.7 for non-conifers. Whole plants in forest products sequestered 7.3 ± 1.0 GtCO₂/year of carbon in 2024, two-thirds of which came from non-conifer species. The carbon-weighted average CP, MR, and CSD of forest products were 43.8 ± 7.2, 18.4 ± 1.6, and 31.1 ± 3.6 years, respectively.

### 4. Other carbons in 2024

The primarily food-oriented use of agricultural products extends plant carbon in the form of animal biomass and their waste, at levels of approximately 18% and 9% by weight, respectively. The lifetime of animal carbon before its release into the atmosphere through respiration ranges from one week in the liver to six weeks in hair (Tieszen et al., 1983). Fecal matter in the form of compost or manure is mineralized in the environment, with a carbon lifetime estimated at a few months.

With 126 MtCO_2_ in 2024, the variations in carbon stocks of aquatic products (fishing and aquaculture) and of animal populations on land (humanity and livestock) were 10 times lower than those of the three main terrestrial plant productions.

### 5. Carbon from cultivated plants in 2024

The carbon involved in harvesting cultivated plants, denoted Har, came from 40.2 ± 1.1 GtCO_2_/year removed from the atmosphere in 2024 (Table 1) for a weighted average CSD half-life of 10.7 ± 2.9 years. The share of annual plants in this total was 27% for a CSD of 6 years.

**Table 1:** Quantities of CO_2_ mobilized by harvesting whole cultivated plants (GtCO_2_/year) in 2024 and weighted average duration of their capture and restitution. SD = standard deviation.

|  | Harvest<br>GtCO<sub>2</sub>/yr | SD | CP<br>yr | RM<br>yr | CDS<br>yr | SD |
| --- | --- | --- | --- | --- | --- | --- |
| Crops | 14.5 | 0.3 | 4.9 | 20.6 | 12.7 | 0.6 |
| Forage | 18.2 | 1.7 | 1.2 | 0.7 | 0.9 | 0.1 |
| Forestry | 7.3 | 1.0 | 43.8 | 18.5 | 31.2 | 3.6 |
| Other | 0.1 | 0.1 | 2.3 | 39.6 | 4.6 | 0.7 |
| Total/Average | 40.2 | 1.1 | 10.3 | 11.2 | 10.7 | 2.9 |

### 6. Net balance of cultivated plants in 2024

The matrix calculation calibrated using the 2024 parameters from Table 2 gives an AW capture of 41.01±1.1 GtCO_2_/year according to (7) and an EW restitution of 37.8 ± 1.8 according to (8).

**Table 2:**
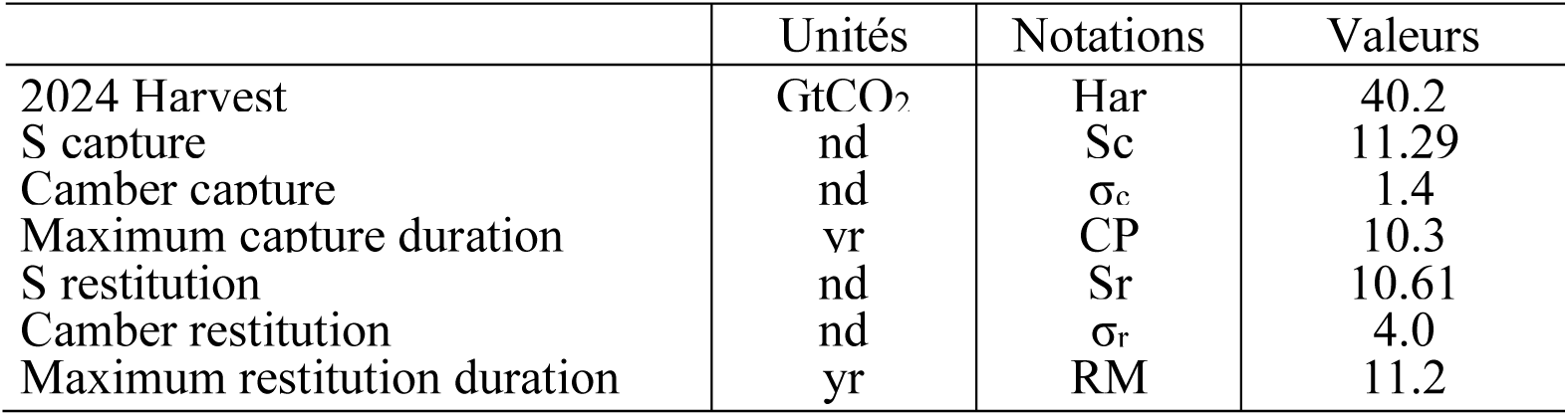
Simulation parameters of the variation of carbon uptake and restitution over time in 2024 (nd: dimensionless).

|  | Unités | Notations | Valeurs |
| --- | --- | --- | --- |
| 2024 Harvest | GtCO<sub>2</sub> | Har | 40.2 |
| S capture | nd | Sc | 11.29 |
| Camber capture | nd | σ<sub>c</sub> | 1.4 |
| Maximum capture duration | yr | CP | 10.3 |
| S restitution | nd | Sr | 10.61 |
| Camber restitution | nd | σ<sub>r</sub> | 4.0 |
| Maximum restitution duration | yr | RM | 11.2 |

The difference between these values and the harvest Har is due to captures and restitutions in years other than 2024 already or still in progress. The net balance of cultivated plants was a sink of 3.2±0.7 GtCO_2_/year in 2024 which enriches the planet’s plant-based carbon stock.

### 7. Anabolism and catabolism of the 2024 harvest

Figure 2 shows the probable chronology of negative carbon capture and positive carbon restitution flow rates mobilized by the 2024 global harvest. Numerically, this is the column vector of the matrix [C(t,n) – R(t,n)]. This chronology is asymmetrical around the harvest year, with the carbon capture (anabolism) rate peaking at -7.1 tCO_2_/yr^2^ and the restitution (catabolism) rate being slightly lower (<6.6 tCO_2_/yr^2^).

**Figure 2:**
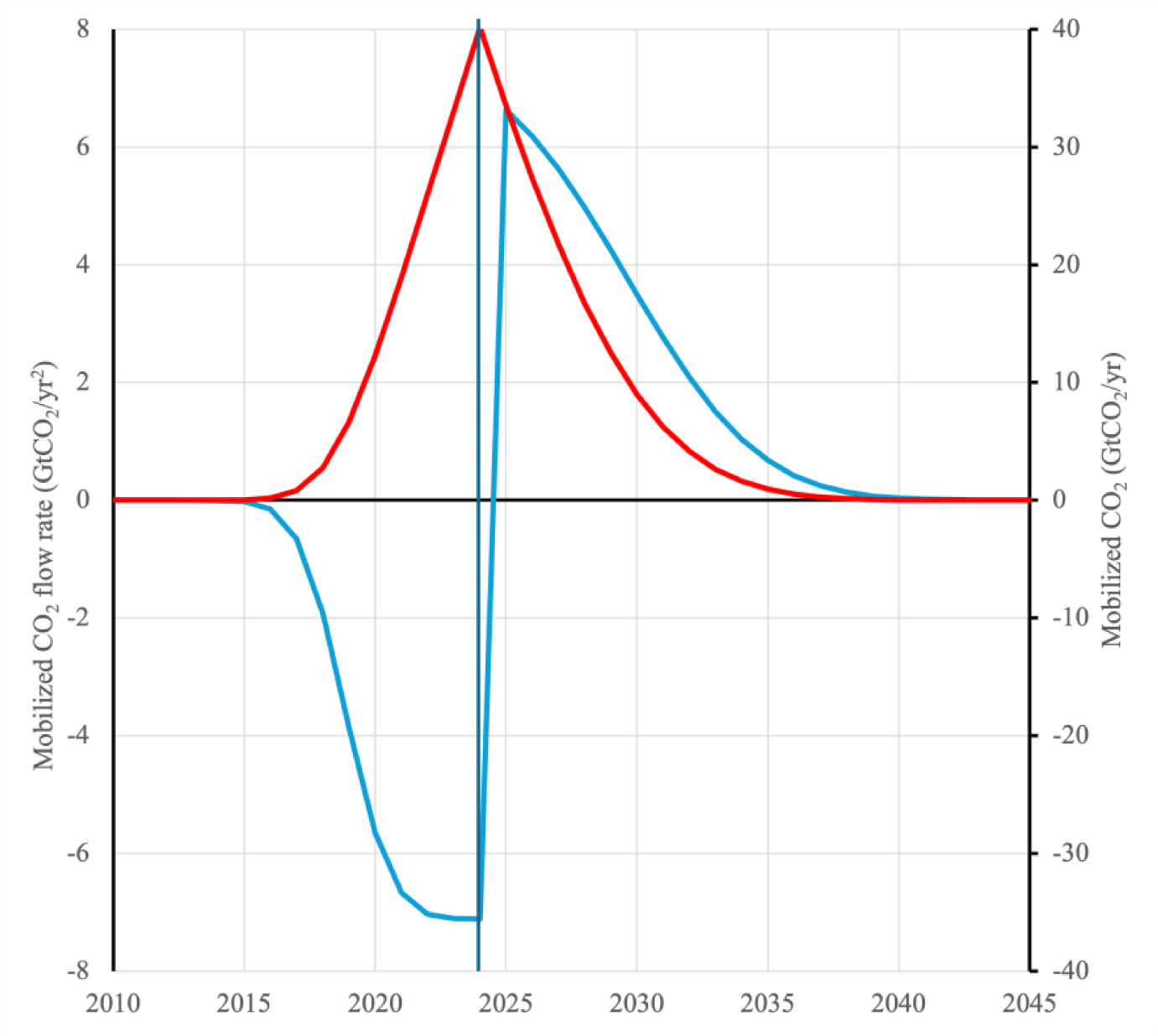
Chronology of CO_2_ mobilization involved in the 2024 harvest of cultivated plants: - in blue, left axis: flow rate of CO_2_ mobilization (GtCO_2_/yr^2^), negative from the atmosphere to plants, and positive in the opposite direction.
- in red on the right axis: CO_2_ mobilized (GtCO_2_/year) with anabolism before harvest and catabolism after.

The integration according to equation (10) of the flow rate absolute values represents the variation of the CO_2_ mobilized by the 2024 harvest. The total duration of anabolism is 12 years and that of catabolism is 23 years. The retention of atmospheric CO_2_ for more than three decades makes cultivated plants a component of carbon budgets.

### 8. Carbon from 1961 to 2024

FAO statistics describe the products of crops, forage, and forests from 1961 onwards. The CO_2_ mobilization of harvests, denoted Har, over the past half-century is determined for each decade and then interpolated between these years. This arrangement is acceptable given the near-linear increase in this mobilization (39% per year, R² = 0.98). The AW capture and EW restitution are deduced from the Har harvests obtained by applying formulas (7) and (8) according to the parameters in Table 2. It was necessary to go back to 1961 to include all stocks restituted in 1970, and to look ahead to 2030 to include all stocks captured in 2024. A boundary effect makes the AW and EW values suspect before 1970; therefore, we only consider the net balances of cultivated plants from that date onward.

Global emissions from fossil fuel combustion (EFOS) reached 38.4 ± 2.3 GtCO_2_/year in 2024, an increase despite a decline in 2020 during the health crisis, according to the Global Change Data Lab (Ritchie et al., 2023). These emissions include those from agriculture and forestry, as well as “other greenhouse gas emissions, the energy mix, and other indicators of potential interest.”

The variation in atmospheric CO_2_ concentration (TAC) measured by Keeling (2001) at the Mauna Loa Observatory (MLO) since 1958 shows a clear upward trend (Figure 3) of more than 2 ppm/year and exhibits annual oscillations with an average increase of 6 ppm from August to April (northern cold season) and an average decrease of 4 ppm from April to August (northern warm season). These oscillations indicate a strong influence from the continents of the Northern Hemisphere, which emit and absorb the largest quantities of CO_2_.

**Figure 3:**
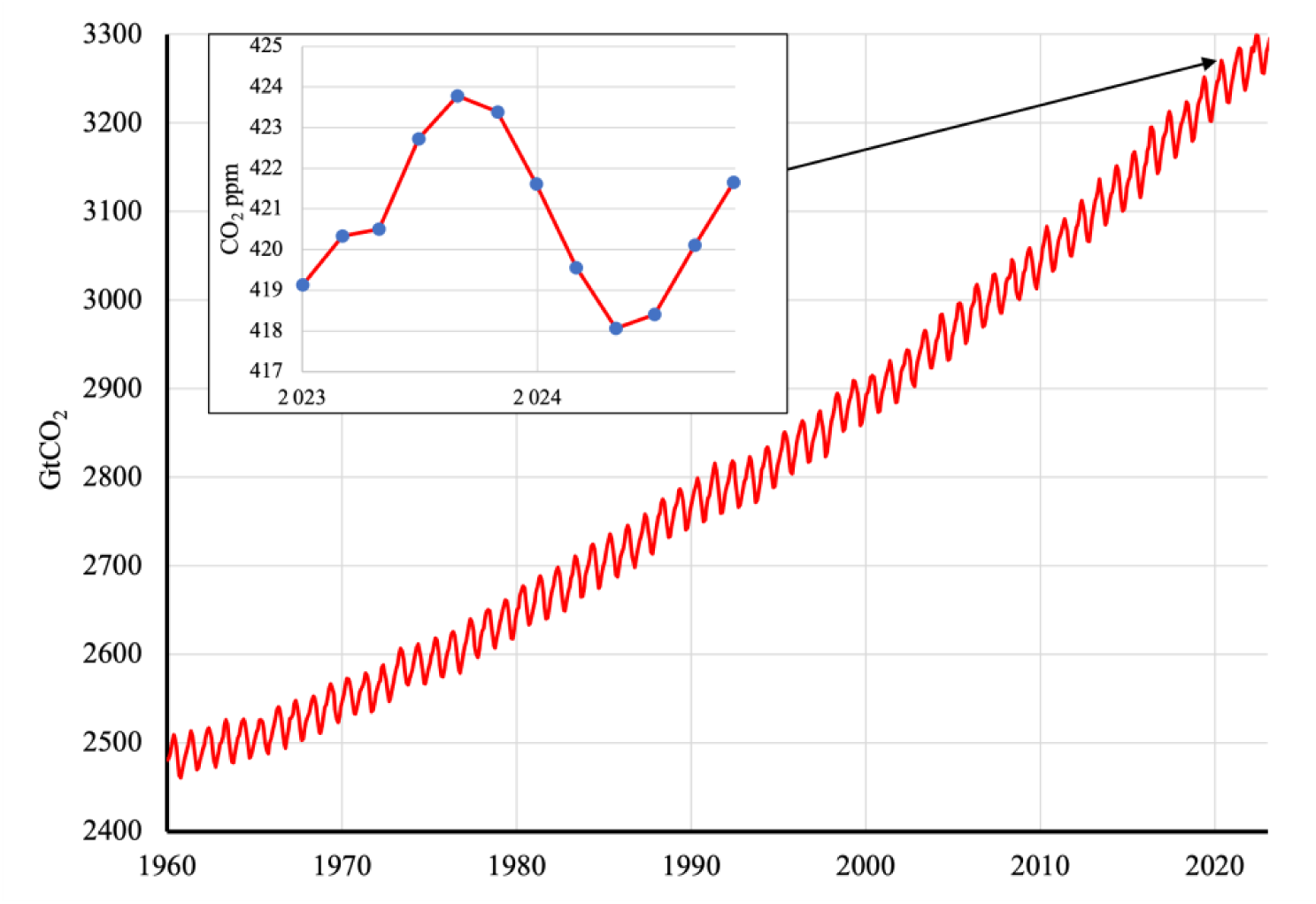
Atmospheric mass of CO_2_ in GtCO_2_, with details for the year 2024 in ppm, based on MLO observations.

The ACC recordings allow us to calculate the variation in mass ACV expressed in GtCO_2_/yr, which is the CO_2_ atmospheric sink.

### 9. Population-driven hypothesis

The population has increased from 3.0 billion capita (Gc) in 1960 to 8.2 Gc in 2024. Figure 4 shows the variation of carbon budget elements relative to the human population size from 1960 to 2024, and their regression lines. These three indices have increased slightly over the last half-century, with shallow slopes indicating their strong proportionality to the population.

**Figure 4:**
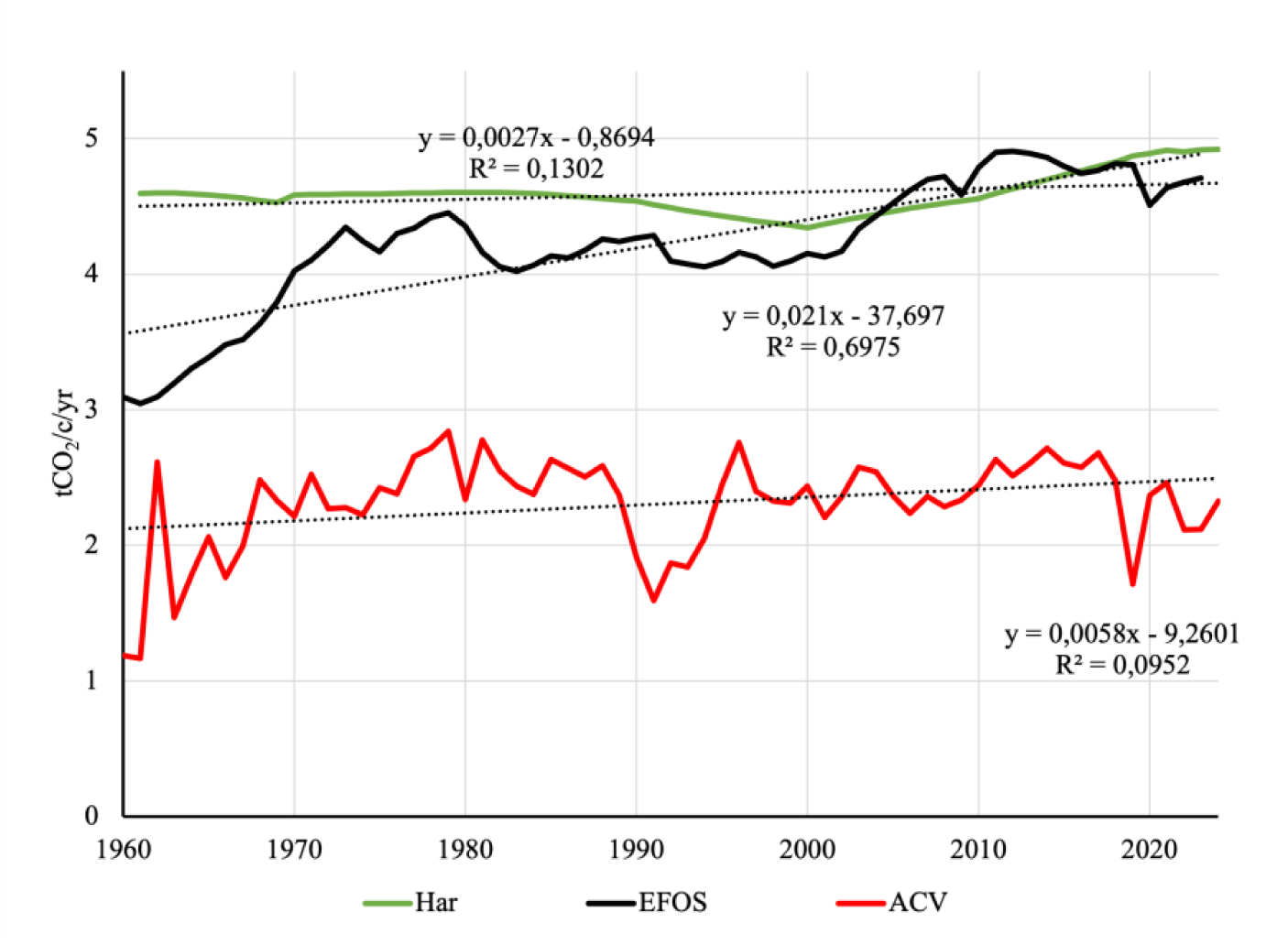
Trends between 1960 and 2024 in cultivated plants harvests (Har), fossil fuel emissions (EFOS), and the atmospheric carbon sink (ACV), per capita of world population (tCO_2_/c/yr).

Fossil fuel emissions (EFOS) relative to the population are increasing by 2.1% per year to over 4.5 tCO_2_ per capita per year (tCO_2_/c/yr) in 2024. This appears to be linked to strong demand from emerging countries, particularly for electricity generation. The upward trend in the atmospheric carbon sink ACV (0.6% per year) seems to be related to that of these fossil fuel emissions. Cultivated plants production Har has been increasing slightly by 0.3% per year to 4.5 tCO_2_/c/yr since 1961. This near-stability could indicate improved demand coverage, particularly related to supply distribution. However, the rate of undernutrition remains high, and progress is needed.

### 10. Carbon budget projections to the end of the current century

According to the UN median scenario (United Nations, 2024) as of July 2024, the population is expected to peak at 10.3 ± 0.81 Gc around 2084. We have projected the evolution of CO_2_ exchanges with the atmosphere until 2100 by adjusting the three variables Har, EFOS, and ACV by the global population size according to this median scenario. This is a provisionally acceptable assumption that minimizes energy projections if the upward trend of this demand continues.

The EFOS source, the ACV sink, and the net AW – EW balance of cultivated plants allow us to specify the contribution of oceans and unexploited continental land areas (OS+) calculated according to equation (12), which completes the carbon budget (Figure 5).

**Figure 5:**
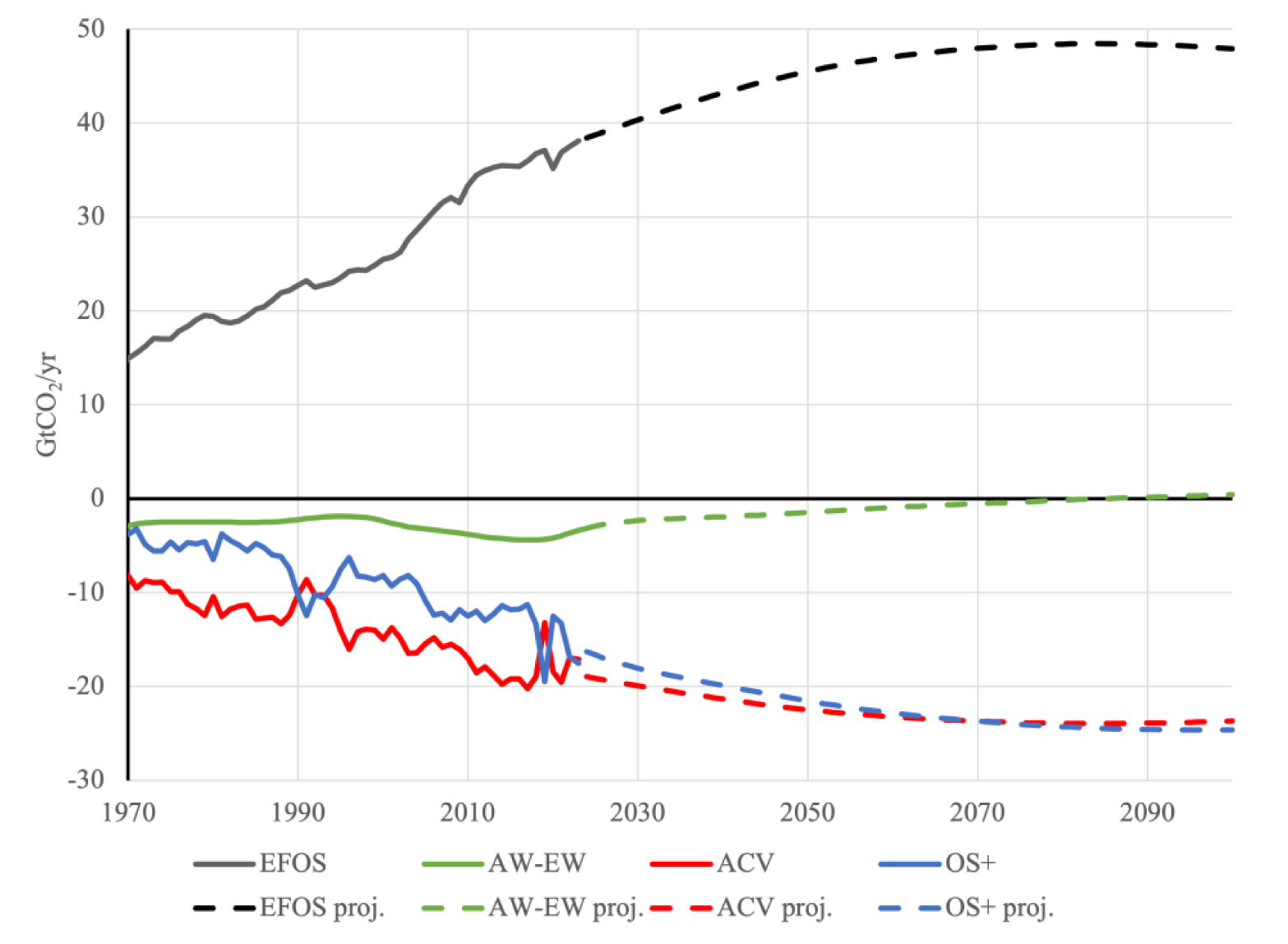
Changes in the four elements of the CO_2_ exchange budget over the last half-century (solid lines) and projections to the end of the century (dashed lines) based on global population growth according to the UN median scenario. EFOS: fossil fuel emissions; AW-EW: net balance of cultivated plants; ACV: mass change in atmospheric CO_2_; OS+: ocean and uncultivated land. Sinks are negative and sources positive. Their algebraic sum is nil.

The EFOS source and the ACV sink continue to grow until the population peak projected for 2084, before declining. The net balance of cultivated plants AW-EW remains a weakening sink until this time before becoming a source. The OS+ sink continues to grow until a peak towards the end of the century.

### 11. Carbon stocks from cultivated plants

The 3D representation (Figure 6) describes the mobilizations by successive decennial harvests represented by the colored bands along the Stocks axis.

**Figure 6:**
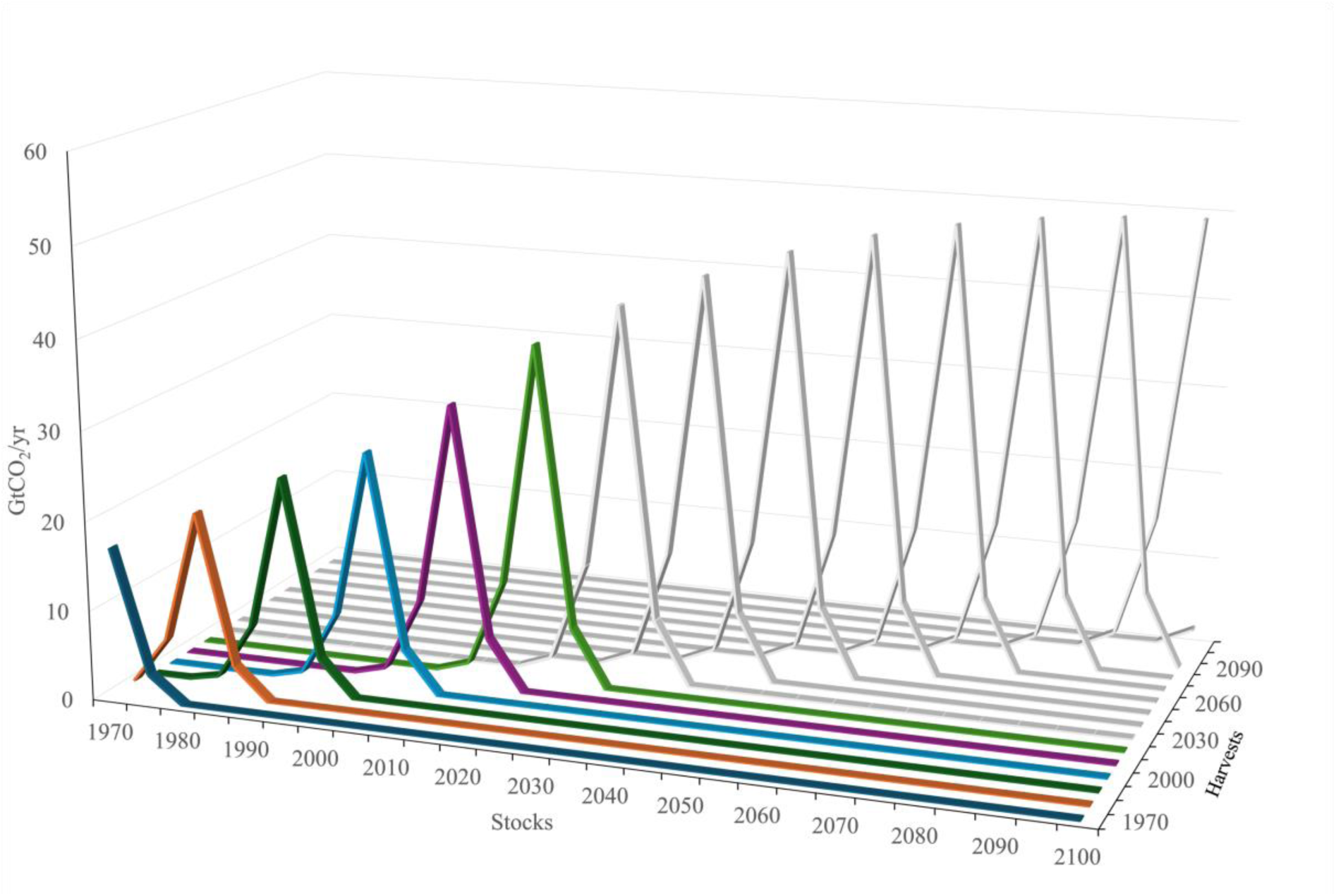
Quantities of atmospheric CO_2_ (GtCO_2_/year) mobilized by cultured plants worldwide for every decade from 1970 to 2020 (in color), and until the end of the century (in gray), as projected according to the UN median scenario for the 2024 human population projection.

The sum of the net balances of crops AW – EW represents the carbon stock contained in vegetation, buildings, stores, shelves, soils, and fauna. Numerically, it is the cumulative sum, since 1970 of the row vectors of the [C+R] matrix according to formula (10). Without considering stocks already built up before 1970, they have increased by 43.4 GtC in 2024 at a rate of 71% per year (R^2^ = 0.98). According to our projections, they will increase by a further 21 GtC, culminating at the horizon of maximum population before declining. These stocks do not include those on unexploited continental land surfaces, which are not included in the statistics and are part of the expanded oceanic carbon sink OS+.

## 4. Discussion

The purpose of this exercise is to better understand the behavior of atmospheric CO₂ by attempting to quantify its variations. It is primarily motivated by the hypothesis that this gas influences tropospheric temperatures. However, some studies (e.g., Richet P., 2021; Koutsoyiannis D. et al., 2023) suggest that the causal link is reversed, meaning that the fate of CO_2_ is influenced by temperature, which has other origins. Even if the hypothesis of the influence of the atmospheric carbon on climate is not verified, it remains useful to understand how it behaves due to its vital importance, especially as it tends to become scarse on a geological timescale.

The results obtained here differ significantly from those published previously (Muller-Feuga, 2025b). The main innovation lies in the integration of the CO_2_ mobilization rate to define the quantities mobilized and their timing. The calculation of the restitution EW, which was significantly underestimated previously, is here equal to the sum of the row vectors of the restitution matrix [R] according to formula (8), after ensuring that the sums of the column vectors of the two matrices [R] and [C] are indeed equal to the harvest Har. These corrections make the ocean and unexploited continental surfaces a sink, which is consistent with the literature. Our projection shows that the net balance of cultivated plants is a vanishing sink until the population maximum before becoming a source. It would continue to build up a significant carbon stock until this horizon.

Apart from atmospheric variation ACV and fossil fuel emissions EFOS, which are accurately measured and provided by public statistics, the figures obtained rather differ from those of the literature. These differences are partly due to the failure of the cited authors to consider cultivated annual plants, partly also to the imprecision of the elements of the budget deduced from the field measurements interpolated and extended to remotely sensed areas, and finally partly since we have not quantified the share of unexploited continental areas.

### 12. Consideration of annual plants

The stock-to-use ratio of 30% of cereals (FAO, 2018) suggests that global stocks are replenished approximately every three years, thus helping to control prices. This is an average with notable exceptions. For example, Japan recently opened its strategic rice reserves, which had been created thirty years prior (The Mainichi, 2025). More generally, the global consumer price index for rice shows a persistent upward trend, encouraging countries to import and build up food reserves. Replenishing these reserves is a strategic imperative for coping with crises in a context of population growth, the end of which is still far off.

We found only one publication in the literature describing the contribution of annual plants to global carbon sequestration (Wolf et al., 2015). Based on the same sources as ours (FAO n.d.), a comparable contribution to our results is mentioned. The orders of magnitude are three times greater than those of authors who do not consider annual plants (e.g. Le Quéré, 2013). However, Wolf et al. (2015) considered that the storage duration did not exceed one year, with restitution through respiration immediately following harvest, which is consistent with the provision excluding annual plants.

This provision only reflects reality in the very rare situation where the harvest is burned. Even in this extreme situation, the underground parts, which represent 30% of the biomass, persist for several years or even decades. The share of annual plants in carbon mobilization was 75% of crops and 27% of all plants produced in 2024, with a carbon retention half-life of 6.0 years. This duration does not justify excluding annual plants, which significantly skews the carbon budget.

### 13. Imprecision of the budgets

Unlike our approach, which relies on FAO trade product statistics, most authors construct carbon emissions from field and satellite surveys, subsequently extended by interpolation models to the global areas thus characterized. It appears that this method runs up against problems of density and precision in the measurements characterizing each vegetation cover, rather than the areas of these covers, which are precisely defined thanks to satellite imagery. Estimates of continental and oceanic absorption and emission vary, fueling controversies, particularly regarding the contribution of primary forests (e.g. Luyssaert et al., 2008; Gundersen et al., 2021; Luyssaert et al., 2021).

Estimates of soil carbon stocks (e.g. Raith et al., 2002; Lal R., 2008; Chen S. et al., 2013; Nissan et al., 2023) mention 1500 to 2400 GtC, much more than the atmosphere (900 GtC) or terrestrial vegetation (450–650 GtC). One-fifth of atmospheric CO_2_ would originate from soils (400 GtCO_2_/yr), making them the main global source, with approximately ten times the emissions from fossil fuels. These estimates should be reconsidered because they assume an increase in atmosphere of 51 ppm/yr, whereas it was only 2.6 ppm/yr in 2024.

### 14. Unexploited Areas

Various attempts have been made to determine the contribution of unexploited vegetation cover to carbon exchange with the atmosphere. This includes old-growth forests, tundras, marshes, peatlands, savannas, scrubland, mangroves, lakes, and rivers. This contribution is not expected to increase with the world population and should even decrease due to urbanization and the expansion of agricultural land through the draining of marshes, deforestation, terrace farming, etc. However, the increase of ACC, which improves crop yields, could offset this trend.

The use of satellite data provides some clarification on the areas in question. For example, Chen C. et al. (2019) estimate the Leaf Area Index (LAI) mix at 17.85 for arable land, 16.72 for forests, 11.5 for other woody vegetation, and 7.85 for pasture. Unexploited vegetation cover would therefore account for 21.3% of the total leaf area. For its part, the FAO estimates agricultural and forest areas at 29% and unexploited land (Other Land) at 13% of the Earth’s land surface in 2023. These differences could be due to the difficulty of distinguishing unexploited from exploited areas, particularly when it comes from pastures of extensive livestocks.

### 15. Area Yields

The area criterion is incomplete for assessing the contribution of these land covers because it does not consider the flux, in other words, the area yield of atmospheric carbon capture, which depends on climatic and edaphic factors, including water, fertilizer, and labor inputs. According to Saugier et al. (2001), the area flux of carbon capture ranges from 3.7 tCO_2_ per ha and per year (/ha/yr) for deserts to 43.1 tCO_2_/ha/yr for tropical forests, with only 10.1 tCO_2_/ha/yr for crops. More recently, Basher and Ackter (2022) observed that forest ecosystems are the main terrestrial carbon sinks.

Based on 2024 FAO statistics, the average area yield weighted by the dry weight of whole cultivated plants worldwide was 8.9 tCO_2_/ha/yr for crops, 5.3 tCO_2_/ha/yr for pastures, and 1.8 tCO_2_/ha/yr for forests. As examples corroborating these figures, forestry in Brazil has adopted a standard post-harvest recovery rate of 0.9 tCO_2_/ha/yr (Vidal et al., 2020), which is half the global yield of forests. The average carbon sequestration of French forests is 3.3 tCO_2_/ha/yr (IGN, 2013), less than that of pastures and crops worldwide. The contribution of forests was significantly overestimated and that of crops significantly underestimated by most authors.

It should be noted here that logging and forest fires do not lead to a loss of capacity to capture and store CO_2_. The soil carbon stock remains in place, while spontaneous or replanted regrowth continues to capture CO_2_ before new logging occurs 25 to 120 years later. If the forest is replaced by pastures or crops, the land it frees up captures CO_2_ at levels more than 10 times higher. Sugarcane and palm oil crops come out on top with more than 40 tCO_2_/ha/yr followed by vegetable and cereal crops with more than 10 tCO_2_/ha/yr.

### 16. The share of unexploited land

The difference between our OS+ sink and estimates of the ocean sink could be the net balance of wild vegetation on unexploited continental land. Our OS+ sink (16.3 ± 2.7 GtCO_2_/yr in 2024) is significantly larger than that attributed to the ocean in the literature, which results from numerous interpolated measurements of CO_2_ and total inorganic carbon fugacity. These measurements would show that the ocean absorbs a quarter of anthropogenic emissions (NOAA, n.d.), or about 10 GtCO_2_/yr. Other estimates of the ocean sink are 12.5 ± 1.5 GtCO_2_/yr (Friedlingstein et al., 2025). Mignot et al. (2025) analyzed the factors influencing the variability of the ocean sink and found that it has been increasing since 1990 with the rise in ACC. It is projected to reach 13.75 ± 0.5 GtCO_2_/yr by 2024, an increase rate of 1.3 ± 0.1 GtCO_2_/yr. This leaves a difference of 3.15 GtCO_2_/year with our OS+ sink, which could represent the net balance of wild vegetation on unexploited continental land. This net balance would then equal our net balance of harvested vegetation (AW-EW = 3.16 ± 0.7 GtCO_2_/yr in 2024).

Another approach to determining the share of unexploited land involves subtracting the net balance of harvested land from the total net balance of vegetation to identify the share of wild vegetation. The total net carbon balance of vegetation in the literature, denoted SLAND-ELUC, is established using two harmonized approaches (Grassi et al., 2023): one based on accounting models partially derived from FAO statistics, and the other on dynamic models, in accordance with the guidelines for researchers (IPCC, 2006; 2019), i.e., excluding annual crops.

Jia et al. (2025) estimated the continental net carbon balance of all plants at 5.5 ± 2.9 GtCO_2_/yr on average between 1992 and 2020. Pan et al. (2024) estimated the net carbon sink constituted by global continental vegetation at 5.9 ± 1.8 GtCO_2_/yr on average during the period 2010–2019, which is of the same order of magnitude. These figures would attribute a share of 53% and 57%, respectively, to cultivated plants among all vegetation. Given that the net carbon balance for plants according to Friedslingstein et al. (2024, 2025) was 4.8 ± 2.9 GtCO_2_/yr in 2024, 1.64 GtCO_2_/yr would remain for uncultivated land in 2024. The distribution would then be one-third for wild plants and two-thirds for cultivated plants among continental vegetation.

The difference between the two approaches could be due to the exclusion of annual plants, which reduces the assessments of the second approach and favors the first, with the same proportion applied to both wild and cultivated plants. However, these figures are within the confidence interval and should be interpreted with caution. In any case, continental vegetation in general, and cultivated vegetation in particular, must be given its due consideration in carbon budgets.

## 5. Conclusion

Analysis of global crop production data published by the FAO has established that these crops capture and store 40.2 ± 1.1 GtCO_2_/yr over a weighted average half-life of 10.7 ± 2.9 years in 2024. The characteristics of these crops have allowed us to calibrate the probabilistic laws governing the temporal distribution of carbon from cultivated plants since 1961.

To project carbon budgets to the end of the century, we assume that fossil fuel emissions, crop production, and the increase in atmospheric carbon dioxide will evolve proportionally to global population growth. This allows us to identify the share of ocean and uncultivated land that will peak at the end of the century. Fossil fuel emissions and carbon stocks from cultivated plants will reach a maximum at the same time as the human population before declining. Simultaneously, the carbon sink provided by cultivated plants would be depleted and become a carbon source.

This analysis, which requires further refinement and expansion, presents an updated view of the role of plants in the global carbon budget. Given the strong correlation between the components of the budget and the world population, projections based on demographics could well describe the major trends of the end of this century.

## 6. Acknowledgements

We are particularly grateful to all the data collectors at FAO, GCDL, NOAA, BP, UN, etc., without whom this analysis would not have been possible. We thank the many people involved in collecting and making global data available and recognize the considerable personnel and time resources devoted to collecting and formatting this invaluable data. We also extend our gratitude to the engineers who wrote the programs that process this data, put it online, and make it easily accessible to users.

## 8. Appendix 1: Cultivated Plants

Fresh weight of global agricultural crop production in 2024 (FAO, n.d.), water content, annual or perennial plant, dry weight, CO_2_ mobilized by whole plant (WP), maximum capture duration (CP), maximum mineralization restitution duration (MR), carbon half-life (CSD). Water content data are from several sources, including https://feedtables.com and https://www.lanutrition.fr.

| Product | Fresh weight t | Water content % | Annual | Dry weight Mt | MtCO<sub>2</sub> WP | CP yr | RM yr | DSC yr |
| --- | --- | --- | --- | --- | --- | --- | --- | --- |
| Sugar cane | 1939.8 | 75 | FALSE | 484.9 | 1486.4 | 5.0 | 0.1 | 2.6 |
| Maize (corn) | 1218.2 | 13.7 | TRUE | 1051.3 | 2817.9 | 0.8 | 10.2 | 5.5 |
| Rice | 820.2 | 12 | TRUE | 721.8 | 1934.7 | 0.8 | 14.5 | 7.6 |
| Wheat | 798.5 | 13.1 | TRUE | 693.9 | 1859.8 | 0.8 | 11.5 | 6.2 |
| Oil palm fruit | 418.7 | 5 | FALSE | 397.8 | 1219.2 | 30.0 | 143.7 | 86.8 |
| Soya beans | 397.7 | 12 | TRUE | 350.0 | 938.0 | 0.8 | 14.5 | 7.6 |
| Potatoes | 390.4 | 78.3 | TRUE | 84.7 | 227.1 | 1.0 | 0.1 | 0.6 |
| Cassava. fresh | 341.9 | 60 | TRUE | 136.8 | 366.6 | 2.0 | 0.2 | 1.1 |
| Sugar beet | 293.6 | 75 | TRUE | 73.4 | 196.8 | 0.8 | 0.1 | 0.5 |
| Other vegetables. fresh n.e.c. | 281.5 | 85 | TRUE | 42.2 | 113.2 | 0.8 | 0.1 | 0.4 |
| Tomatoes | 188.5 | 94 | TRUE | 11.3 | 30.3 | 0.8 | 0.1 | 0.4 |
| Barley | 142.0 | 12.8 | TRUE | 123.8 | 331.9 | 0.8 | 12.2 | 6.5 |
| Bananas | 139.4 | 75.8 | FALSE | 33.7 | 103.4 | 3.0 | 0.1 | 1.6 |
| Onions and shallots. dry (excluding dehydrated) | 108.3 | 20 | TRUE | 86.6 | 232.1 | 0.8 | 3.8 | 2.3 |
| Watermelons | 105.0 | 90 | TRUE | 10.5 | 28.1 | 0.8 | 0.1 | 0.4 |
| Apples | 97.9 | 85 | FALSE | 14.7 | 45.0 | 1.0 | 0.1 | 0.5 |
| Yams | 92.4 | 75 | FALSE | 23.1 | 70.8 | 0.8 | 0.1 | 0.5 |
| Sweet potatoes | 90.9 | 75 | FALSE | 22.7 | 69.7 | 0.8 | 0.1 | 0.5 |
| Rape or colza seed | 87.9 | 12 | TRUE | 77.3 | 207.2 | 0.8 | 14.5 | 7.6 |
| Cucumbers and gherkins | 87.8 | 80 | TRUE | 17.6 | 47.1 | 0.8 | 0.1 | 0.5 |
| Grapes | 75.9 | 81 | FALSE | 14.4 | 44.2 | 20.0 | 0.1 | 10.0 |
| Cabbages | 75.5 | 93 | TRUE | 5.3 | 14.2 | 0.8 | 0.1 | 0.4 |
| Seed cotton. unginned | 72.8 | 20 | TRUE | 58.3 | 156.1 | 10.0 | 3.8 | 6.9 |
| Oranges | 67.2 | 90 | FALSE | 6.7 | 20.6 | 20.0 | 0.1 | 10.0 |
| Coconuts. in shell | 65.5 | 45 | FALSE | 36.0 | 110.5 | 20.0 | 0.5 | 10.2 |
| Sorghum | 64.3 | 12.2 | TRUE | 56.4 | 151.2 | 0.8 | 13.9 | 7.3 |
| Mangoes. guavas and mangosteens | 62.2 | 83 | FALSE | 10.6 | 32.4 | 20.0 | 0.1 | 10.0 |
| Groundnuts. excluding shelled | 57.6 | 45 | TRUE | 31.7 | 84.9 | 3.0 | 0.5 | 1.7 |
| Eggplants (aubergines) | 57.5 | 80 | TRUE | 11.5 | 30.8 | 0.8 | 0.1 | 0.5 |
| Sunflower seed | 52.2 | 7.2 | TRUE | 48.5 | 129.9 | 0.8 | 55.3 | 28.1 |
| Tangerines. mandarins. clementines | 51.9 | 90 | FALSE | 5.2 | 15.9 | 1.0 | 0.1 | 0.5 |
| Mushrooms and truffles | 51.6 | 75 | TRUE | 12.9 | 34.6 | 1.0 | 0.1 | 0.6 |
| Carrots and turnips | 45.1 | 80 | TRUE | 9.0 | 24.2 | 0.8 | 0.1 | 0.5 |
| Chillies and peppers. green (Capsicum spp. and Pimenta) | 44.8 | 60 | TRUE | 17.9 | 48.0 | 0.8 | 0.2 | 0.5 |
| Plantains and cooking bananas | 44.6 | 75 | FALSE | 11.1 | 34.2 | 5.0 | 0.1 | 2.6 |
| Other fruits. n.e.c. | 39.2 | 80 | FALSE | 7.8 | 24.0 | 3.0 | 0.1 | 1.6 |
| Pumpkins. squash and gourds | 34.6 | 12 | TRUE | 30.4 | 81.6 | 0.8 | 14.5 | 7.6 |
| Tea leaves | 32.7 | 10.5 | FALSE | 29.3 | 89.8 | 50.0 | 20.6 | 35.3 |
| Millet | 30.9 | 10.4 | TRUE | 27.6 | 74.1 | 0.8 | 21.1 | 10.9 |
| Spinach | 30.7 | 60 | TRUE | 12.3 | 32.9 | 1.0 | 0.2 | 0.6 |
| Beans. dry | 30.3 | 10.11 | TRUE | 27.2 | 72.9 | 0.8 | 22.7 | 11.8 |
| Green garlic | 29.7 | 80 | TRUE | 5.9 | 15.9 | 0.8 | 0.1 | 0.5 |
| Pineapples | 29.4 | 75 | TRUE | 7.3 | 19.7 | 0.8 | 0.1 | 0.5 |
| Cantaloupes and other melons | 28.2 | 90 | TRUE | 2.8 | 7.6 | 1.0 | 0.1 | 0.5 |
| Lettuce and chicory | 28.2 | 80 | TRUE | 5.6 | 15.1 | 0.8 | 0.1 | 0.5 |
| Peaches and nectarines | 27.9 | 90 | FALSE | 2.8 | 8.5 | 20.0 | 0.1 | 10.0 |
| Pears | 27.6 | 84 | FALSE | 4.4 | 13.6 | 20.0 | 0.1 | 10.0 |
| Cauliflowers and broccoli | 26.9 | 90 | TRUE | 2.7 | 7.2 | 1.0 | 0.1 | 0.5 |
| Other beans. green | 25.9 | 45 | TRUE | 14.3 | 38.2 | 0.8 | 0.5 | 0.6 |
| Olives | 25.6 | 60 | FALSE | 10.2 | 31.3 | 20.0 | 0.2 | 10.1 |
| Other tropical fruits. n.e.c. | 23.9 | 80 | FALSE | 4.8 | 14.7 | 0.8 | 0.1 | 0.5 |
| Lemons and limes | 23.2 | 90 | FALSE | 2.3 | 7.1 | 0.8 | 0.1 | 0.4 |
| Oats | 22.4 | 12.4 | TRUE | 19.7 | 52.7 | 0.8 | 13.3 | 7.1 |
| Peas. green | 21.7 | 75 | TRUE | 5.4 | 14.6 | 0.8 | 0.1 | 0.5 |
| Taro | 18.2 | 75 | TRUE | 4.6 | 12.2 | 0.8 | 0.1 | 0.5 |
| Chick peas. dry | 16.9 | 12.7 | TRUE | 14.7 | 39.5 | 0.8 | 12.5 | 6.7 |
| Natural rubber in primary forms | 14.8 | 20 | FALSE | 11.9 | 36.4 | 30.0 | 3.8 | 16.9 |
| Papayas | 14.7 | 89.4 | FALSE | 1.6 | 4.8 | 20.0 | 0.1 | 10.0 |
| Peas. dry | 14.2 | 12 | TRUE | 12.5 | 33.4 | 0.8 | 14.5 | 7.6 |
| Other citrus fruit. n.e.c. | 13.4 | 90 | FALSE | 1.3 | 4.1 | 20.0 | 0.1 | 10.0 |
| Okra | 13.3 | 24 | TRUE | 10.1 | 27.0 | 20.0 | 2.4 | 11.2 |
| Plums and sloes | 12.7 | 80 | FALSE | 2.5 | 7.8 | 1.0 | 0.1 | 0.6 |
| Triticale | 12.4 | 13.2 | TRUE | 10.8 | 28.9 | 0.8 | 11.3 | 6.0 |
| Rye | 11.6 | 12 | TRUE | 10.2 | 27.4 | 0.8 | 14.5 | 7.6 |
| Coffee. green | 11.3 | 25 | FALSE | 8.5 | 25.9 | 20.0 | 2.1 | 11.1 |
| Avocados | 11.2 | 75 | FALSE | 2.8 | 8.6 | 20.0 | 0.1 | 10.1 |
| Strawberries | 10.7 | 90 | TRUE | 1.1 | 2.9 | 0.8 | 0.1 | 0.4 |
| Cow peas. dry | 10.1 | 12 | TRUE | 8.9 | 23.8 | 0.8 | 14.5 | 7.6 |
| Dates | 9.9 | 75 | FALSE | 2.5 | 7.6 | 20.0 | 0.1 | 10.1 |
| Pomelos and grapefruits | 9.9 | 90 | FALSE | 1.0 | 3.0 | 1.0 | 0.1 | 0.5 |
| Edible roots and tubers with high starch or inulin content. | 9.8 | 78 | TRUE | 2.2 | 5.8 | 0.8 | 0.1 | 0.5 |
| Green corn (maize) | 9.5 | 75 | TRUE | 2.4 | 6.3 | 1.0 | 0.1 | 0.6 |
| Asparagus | 8.9 | 80 | FALSE | 1.8 | 5.5 | 1.0 | 0.1 | 0.6 |
| Cereals n.e.c. | 8.0 | 13.2 | TRUE | 7.0 | 18.7 | 0.8 | 11.3 | 6.0 |
| Lentils. dry | 7.9 | 11.6 | TRUE | 7.0 | 18.7 | 0.8 | 15.8 | 8.3 |
| Sesame seed | 6.7 | 12 | TRUE | 5.9 | 15.8 | 0.8 | 14.5 | 7.6 |
| Broad beans and horse beans. dry | 6.3 | 25 | TRUE | 4.7 | 12.6 | 0.8 | 2.1 | 1.5 |
| Unmanufactured tobacco | 6.0 | 60 | TRUE | 2.4 | 6.5 | 3.0 | 0.2 | 1.6 |
| Onions and shallots. green | 5.6 | 80 | TRUE | 1.1 | 3.0 | 0.8 | 0.1 | 0.5 |
| Chillies and peppers, dry (Capsicum spp., Pimenta spp.). | 5.5 | 25 | TRUE | 4.1 | 11.0 | 0.8 | 2.1 | 1.5 |
| Cocoa beans | 5.2 | 12 | FALSE | 4.6 | 14.1 | 30.0 | 14.5 | 22.2 |
| Persimmons | 5.0 | 80 | FALSE | 1.0 | 3.1 | 20.0 | 0.1 | 10.1 |
| Ginger, raw | 4.9 | 75 | TRUE | 1.2 | 3.3 | 0.8 | 0.1 | 0.5 |
| Pigeon peas. dry | 4.7 | 12 | TRUE | 4.1 | 11.0 | 0.8 | 14.5 | 7.6 |
| Apricots | 4.6 | 80 | FALSE | 0.9 | 2.8 | 20.0 | 0.1 | 10.1 |
| Other pulses n.e.c. | 4.4 | 12 | TRUE | 3.9 | 10.5 | 0.8 | 14.5 | 7.6 |
| Kiwi fruit | 4.4 | 80 | FALSE | 0.9 | 2.7 | 20.0 | 0.1 | 10.1 |
| Walnuts. in shell | 4.3 | 12 | FALSE | 3.8 | 11.7 | 20.0 | 14.5 | 17.2 |
| Cashew nuts. in shell | 4.2 | 45 | FALSE | 2.3 | 7.1 | 20.0 | 0.5 | 10.2 |
| Almonds. in shell | 4.0 | 45 | FALSE | 2.2 | 6.7 | 20.0 | 0.5 | 10.2 |
| Jute, raw or retted | 3.7 | 8 | FALSE | 3.4 | 10.4 | 0.8 | 41.9 | 21.4 |
| Other stimulant. spice and aromatic crops. n.e.c. | 3.3 | 25 | TRUE | 2.5 | 6.7 | 0.8 | 2.1 | 1.5 |
| Cherries | 3.1 | 80 | FALSE | 0.6 | 1.9 | 20.0 | 0.1 | 10.1 |
| Linseed | 3.0 | 8 | TRUE | 2.8 | 7.4 | 0.8 | 41.9 | 21.4 |
| Anise, badian, coriander, cumin, caraway, fennel and | 2.7 | 25 | TRUE | 2.0 | 5.4 | 0.8 | 2.1 | 1.5 |
| Other oil seeds. n.e.c. | 2.4 | 20 | FALSE | 1.9 | 5.9 | 0.8 | 3.8 | 2.3 |
| Areca nuts | 2.3 | 12 | FALSE | 2.1 | 6.3 | 20.0 | 14.5 | 17.2 |
| Leeks and other alliaceous vegetables | 2.1 | 80 | TRUE | 0.4 | 1.1 | 0.8 | 0.1 | 0.5 |
| Chestnuts, in shell | 2.1 | 65 | FALSE | 0.7 | 2.3 | 20.0 | 0.2 | 10.1 |
| Maté leaves | 2.0 | 25 | TRUE | 1.5 | 4.0 | 0.8 | 2.1 | 1.5 |
| Buckwheat | 2.0 | 12 | TRUE | 1.7 | 4.6 | 1.0 | 14.5 | 7.7 |
| Castor oil seeds | 1.8 | 12 | TRUE | 1.6 | 4.3 | 3.0 | 14.5 | 8.7 |
| Coir, raw | 1.8 | 12 | FALSE | 1.6 | 4.8 | 80.0 | 14.5 | 47.2 |
| Artichokes | 1.7 | 80 | TRUE | 0.3 | 0.9 | 1.0 | 0.1 | 0.6 |
| Mixed grain | 1.6 | 12 | TRUE | 1.4 | 3.8 | 0.8 | 14.5 | 7.6 |
| Lupins | 1.5 | 25 | TRUE | 1.1 | 3.0 | 0.8 | 2.1 | 1.5 |
| Sour cherries | 1.5 | 80 | FALSE | 0.3 | 0.9 | 20.0 | 0.1 | 10.1 |
| Broad beans and horse beans. green | 1.5 | 12 | TRUE | 1.3 | 3.4 | 0.8 | 14.5 | 7.6 |
| Blueberries | 1.4 | 90 | FALSE | 0.1 | 0.4 | 3.0 | 0.1 | 1.5 |
| Cashewapple | 1.4 | 12 | FALSE | 1.2 | 3.7 | 30.0 | 14.5 | 22.2 |
| Pistachios. in shell | 1.4 | 12 | FALSE | 1.2 | 3.7 | 30.0 | 14.5 | 22.2 |
| Figs | 1.3 | 12 | FALSE | 1.2 | 3.6 | 20.0 | 14.5 | 17.2 |
| Flax, raw or retted | 1.3 | 8 | TRUE | 1.2 | 3.2 | 0.8 | 41.9 | 21.4 |
| Hazelnuts. in shell | 1.2 | 25 | FALSE | 0.9 | 2.8 | 20.0 | 2.1 | 11.1 |
| String beans | 1.2 | 80 | TRUE | 0.2 | 0.6 | 0.8 | 0.1 | 0.5 |
| Other nuts (excluding wild edible nuts and groundnuts), in | 1.1 | 25 | FALSE | 0.8 | 2.5 | 20.0 | 2.1 | 11.1 |
| Other berries and fruits of the genus vaccinium n.e.c. | 1.1 | 90 | FALSE | 0.1 | 0.3 | 0.8 | 0.1 | 0.4 |
| Tallowtree seeds | 1.1 | 25 | FALSE | 0.8 | 2.4 | 0.8 | 2.1 | 1.5 |
| Melonseed | 1.0 | 25 | TRUE | 0.7 | 2.0 | 0.8 | 2.1 | 1.5 |
| Pepper (Piper spp.), raw | 0.9 | 12 | FALSE | 0.8 | 2.5 | 25.0 | 14.5 | 19.7 |
| Raspberries | 0.9 | 90 | FALSE | 0.1 | 0.3 | 3.0 | 0.1 | 1.5 |
| Other sugar crops n.e.c. | 0.9 | 10 | FALSE | 0.8 | 2.5 | 0.8 | 23.4 | 12.1 |
| Mustard seed | 0.9 | 25 | TRUE | 0.6 | 1.7 | 0.8 | 2.1 | 1.5 |
| Karite nuts (sheanuts) | 0.8 | 25 | FALSE | 0.6 | 1.9 | 20.0 | 2.1 | 11.1 |
| Currants | 0.7 | 80 | FALSE | 0.1 | 0.4 | 3.0 | 0.1 | 1.6 |
| Quinces | 0.7 | 90 | FALSE | 0.1 | 0.2 | 30.0 | 0.1 | 15.0 |
| Safflower seed | 0.7 | 25 | TRUE | 0.5 | 1.4 | 0.8 | 2.1 | 1.5 |
| Fonio | 0.7 | 90 | TRUE | 0.1 | 0.2 | 0.8 | 0.1 | 0.4 |
| Other fibre crops, raw, n.e.c. | 0.7 | 24 | FALSE | 0.5 | 1.5 | 0.8 | 2.4 | 1.6 |
| Vetches | 0.7 | 25 | TRUE | 0.5 | 1.3 | 0.8 | 2.1 | 1.5 |
| Cranberries | 0.6 | 85 | FALSE | 0.1 | 0.3 | 3.0 | 0.1 | 1.5 |
| Other stone fruits | 0.6 | 80 | FALSE | 0.1 | 0.4 | 20.0 | 0.1 | 10.1 |
| Tung nuts | 0.4 | 25 | FALSE | 0.3 | 1.0 | 20.0 | 2.1 | 11.1 |
| Yautia | 0.4 | 93 | TRUE | 0.0 | 0.1 | 0.8 | 0.1 | 0.4 |
| True hemp, raw or retted | 0.4 | 25 | TRUE | 0.3 | 0.7 | 1.0 | 2.1 | 1.6 |
| Kola nuts | 0.3 | 25 | FALSE | 0.2 | 0.7 | 20.0 | 2.1 | 11.1 |
| Canary seed | 0.3 | 12 | TRUE | 0.3 | 0.7 | 0.8 | 14.5 | 7.6 |
| Kapok fruit | 0.3 | 25 | FALSE | 0.2 | 0.7 | 20.0 | 2.1 | 11.1 |
| Cinnamon and cinnamon-tree flowers. raw | 0.2 | 12 | FALSE | 0.2 | 0.7 | 4.0 | 14.5 | 9.2 |
| Bambara beans, dry | 0.2 | 24 | TRUE | 0.2 | 0.5 | 0.8 | 2.4 | 1.6 |
| Sisal, raw | 0.2 | 24 | FALSE | 0.2 | 0.5 | 5.0 | 2.4 | 3.7 |
| Kenaf, and other textile bast fibres, raw or retted | 0.2 | 24 | TRUE | 0.2 | 0.5 | 0.8 | 2.4 | 1.6 |
| Cloves (whole stems). raw | 0.2 | 65 | TRUE | 0.1 | 0.2 | 1.0 | 0.2 | 0.6 |
| Nutmeg, mace, cardamoms, raw | 0.2 | 12 | FALSE | 0.2 | 0.5 | 30.0 | 14.5 | 22.2 |
| Quinoa | 0.2 | 12 | TRUE | 0.1 | 0.4 | 0.8 | 14.5 | 7.6 |
| Other pome fruits | 0.2 | 80 | FALSE | 0.0 | 0.1 | 20.0 | 0.1 | 10.1 |
| Hop cones | 0.2 | 24 | FALSE | 0.1 | 0.4 | 50.0 | 2.4 | 26.2 |
| Abaca, manila hemp, raw | 0.1 | 24 | TRUE | 0.1 | 0.2 | 0.8 | 2.4 | 1.6 |
| Gooseberries | 0.1 | 80 | FALSE | 0.0 | 0.1 | 3.0 | 0.1 | 1.6 |
| Brazil nuts, in shell | 0.1 | 12 | FALSE | 0.1 | 0.2 | 20.0 | 14.5 | 17.2 |
| Ramie, raw or retted | 0.1 | 24 | TRUE | 0.0 | 0.1 | 0.8 | 2.4 | 1.6 |
| Locust beans (carobs) | 0.1 | 80 | FALSE | 0.0 | 0.0 | 20.0 | 0.1 | 10.1 |
| Agave fibres, raw, n.e.c. | 0.0 | 8 | FALSE | 0.0 | 0.1 | 40.0 | 41.9 | 41.0 |
| Peppermint, spearmint | 0.0 | 75 | TRUE | 0.0 | 0.0 | 0.8 | 0.1 | 0.5 |
| Hempseed | 0.0 | 8 | TRUE | 0.0 | 0.1 | 0.8 | 41.9 | 21.4 |
| Chicory roots | 0.0 | 24 | TRUE | 0.0 | 0.0 | 0.8 | 2.4 | 1.6 |
| Poppy seed | 0.0 | 12 | TRUE | 0.0 | 0.0 | 0.8 | 14.5 | 7.6 |
| Pyrethrum, dried flowers | 0.0 | 12 | TRUE | 0.0 | 0.0 | 0.8 | 14.5 | 7.6 |
| Vanilla, raw | 0.0 | 24 | FALSE | 0.0 | 0.0 | 10.0 | 2.4 | 6.2 |
| Jojoba seeds | 0.0 | 12 | TRUE | 0.0 | 0.0 | 3.0 | 14.5 | 8.7 |
|  | 9861 | 46.0 |  | 5228.72 | 14474.14 | 4.9 | 20.6 | 12.7 |

## 9. Appendix 2: Forages

Fresh weight of global livestock products in 2023 (FAO, n.d.), anhydrous forages consumed by these livestock, CO_2_ mobilized, duration of photosynthetic carbon capture (CP), duration of mineralization restitution (MR), and half-life of plant carbon (DSC), ranked in descending order of carbon weight. The conversion rate applied is 4.336 (Mottet et al., 2017). The C/ms ratio of annual plants is used, i.e., 0.425. The resulting carbon capture is reduced by 14% to remove the portion already included in Appendix 1.

| Product | Fresh weight<br>(Mt) | Dry forage<br>(Mt) | Mt CO<sub>2</sub> | CP<br>yr | RM<br>yr | DSC<br>yr |
| --- | --- | --- | --- | --- | --- | --- |
| Raw milk of cattle | 797.3 | 3457.0 | 11151.54 | 1.0 | 0.5 | 0.8 |
| Raw milk of buffalo | 153.0 | 663.6 | 2140.5 | 1 | 0.5 | 0.75 |
| Meat of chickens, fresh or chilled | 128.3 | 556.3 | 1794.6 | 0.8 | 0.5 | 0.65 |
| Meat of pig with the bone, fresh or chilled | 125.4 | 543.5 | 1753.4 | 2 | 0.5 | 1.25 |
| Hen eggs in shell, fresh | 93.7 | 406.3 | 1310.5 | 0.8 | 0.04 | 0.42 |
| Meat of cattle with the bone, fresh or chilled | 69.7 | 302.4 | 975.6 | 3 | 0.5 | 1.75 |
| Raw milk of goats | 20.4 | 88.6 | 285.8 | 1 | 0.5 | 0.75 |
| Meat of sheep, fresh or chilled | 11.5 | 50.0 | 161.4 | 2 | 0.5 | 1.25 |
| Fat of pigs | 11.4 | 49.3 | 159.0 | 0.8 | 10 | 5.4 |
| Raw milk of sheep | 9.8 | 42.6 | 137.3 | 1 | 0.5 | 0.75 |
| Edible offal of cattle, fresh, chilled or frozen | 9.7 | 42.0 | 135.6 | 3 | 0.5 | 1.75 |
| Raw hides and skins of cattle | 8.8 | 38.0 | 122.5 | 0.8 | 10 | 5.4 |
| Edible offal of pigs, fresh, chilled or frozen | 8.2 | 35.6 | 114.8 | 2 | 0.5 | 1.25 |
| Meat of goat, fresh or chilled | 7.7 | 33.6 | 108.3 | 2 | 0.5 | 1.25 |
| Meat of ducks, fresh or chilled | 7.3 | 31.8 | 102.6 | 3 | 0.5 | 1.75 |
| Meat of buffalo, fresh or chilled | 7.3 | 31.6 | 102.1 | 0.8 | 0.5 | 0.65 |
| Eggs from other birds in shell, fresh, n.e.c. | 6.3 | 27.3 | 88.1 | 0.8 | 0.04 | 0.42 |
| Meat of geese, fresh or chilled | 5.4 | 23.4 | 75.6 | 0.8 | 0.5 | 0.65 |
| Meat of turkeys, fresh or chilled | 5.1 | 22.0 | 71.1 | 0.8 | 0.5 | 0.65 |
| Raw milk of camel | 4.2 | 18.2 | 58.6 | 1 | 0.5 | 0.75 |
| Cattle fat, unrendered | 3.4 | 14.8 | 47.7 | 0.8 | 10 | 5.4 |
| Raw hides and skins of sheep or lambs | 2.2 | 9.4 | 30.4 | 0.8 | 10 | 5.4 |
| Game meat, fresh, chilled or frozen | 2.1 | 8.9 | 28.7 | 2 | 0.5 | 1.25 |
| Natural honey | 2.0 | 8.6 | 27.8 | 1 | 0.5 | 0.75 |
| Edible offal of sheep, fresh, chilled or frozen | 2.0 | 8.6 | 27.7 | 0.8 | 10 | 5.4 |
| Shorn wool, greasy, including fleece-washed | 1.7 | 7.4 | 23.7 | 0.8 | 10 | 5.4 |
| Other meat of mammals, fresh or chilled | 1.6 | 6.8 | 21.9 | 0.8 | 0.5 | 0.65 |
| Raw hides and skins of goats or kids | 1.5 | 6.6 | 21.3 | 2 | 0.2 | 1.1 |
| Raw hides and skins of buffaloes | 1.5 | 6.5 | 21.0 | 0.8 | 10 | 5.4 |
| Edible offal of goat, fresh, chilled or frozen | 1.5 | 6.4 | 20.7 | 1 | 0.5 | 0.75 |
| Edible offal of buffalo, fresh, chilled or frozen | 1.2 | 5.4 | 17.5 | 3 | 0.5 | 1.75 |
| Meat of rabbits and hares, fresh or chilled | 0.8 | 3.7 | 11.8 | 4 | 0.5 | 2.25 |
| Horse meat, fresh or chilled | 0.7 | 3.2 | 10.5 | 0.8 | 0.5 | 0.65 |
| Meat of camels, fresh or chilled | 0.7 | 2.9 | 9.2 | 0.8 | 10 | 5.4 |
| Sheep fat, unrendered | 0.6 | 2.8 | 9.0 | 3 | 0.5 | 1.75 |
| Silk-worm cocoons suitable for reeling | 0.5 | 2.0 | 6.5 | 0 | 2 | 1 |
| Buffalo fat, unrendered | 0.4 | 1.9 | 6.2 | 0.8 | 10 | 5.4 |
| Goat fat, unrendered | 0.3 | 1.4 | 4.4 | 0.8 | 10 | 5.4 |
| Edible offals of horses and other equines, fresh, | 0.1 | 0.4 | 1.3 | 3 | 0.5 | 1.75 |
| Edible offals of camels and other camelids, fresh, | 0.1 | 0.4 | 1.3 | 3 | 0.5 | 1.75 |
| Meat of asses, fresh or chilled | 0.1 | 0.3 | 1.0 | 3 | 0.5 | 1.75 |
| Beeswax | 0.1 | 0.3 | 0.9 | 0 | 100 | 50 |
| Meat of other domestic camelids, fresh or chilled | 0.0 | 0.1 | 0.5 | 3 | 0.5 | 1.75 |
| Fat of camels | 0.0 | 0.1 | 0.4 | 0.8 | 10 | 5.4 |
| Snails, fresh, chilled, frozen, dried, salted or in | 0.0 | 0.1 | 0.3 | 0.8 | 0.5 | 0.65 |
| Meat of other domestic rodents, fresh or chilled | 0.0 | 0.1 | 0.3 | 0.8 | 0.5 | 0.65 |
| Meat of pigeons and other birds n.e.c., fresh, | 0.0 | 0.1 | 0.2 | 0.8 | 0.5 | 0.65 |
| Meat of mules, fresh or chilled | 0.0 | 0.0 | 0.1 | 4 | 0.5 | 2.25 |
|  | 1515.8 | 6572.6 | 21201.6 | 1.18 | 0.68 | 0.93 |
| Mottet - 14% |  | 5652.4 | 18233.3 |  |  |  |

## 10. Appendix 3: Forestry Products

Volume of global forest products in 2024 (FAO, n.d.), dry weight, CO_2_ mobilized, growth period and storage (GP), and consumption-mineralization period (CM).

| Item | Mm<sup>3</sup> | Density<br>t/m<sup>3</sup> | Dry<br>Mt | CO<sub>2</sub>/dm | MtCO<sub>2</sub>/yr | CP<br>yr | RM yr | DSC<br>yr |
| --- | --- | --- | --- | --- | --- | --- | --- | --- |
| Wood fuel, non-coniferous | 1711 | 0.70 | 1197 | 1.767 | 3322.7 | 50 | 2 | 26 |
| Sawlogs and veneer logs, coniferous | 706 | 0.6 | 423. | 1.852 | 1232.3 | 30 | 30 | 30 |
| Sawlogs and veneer logs, non-coniferous | 393 | 0.70 | 275. | 1.767 | 763.3 | 50 | 30 | 40 |
| Pulpwood, round and split, non-coniferous | 384 | 0.70 | 268. | 1.767 | 746.2 | 50 | 30 | 40 |
| Pulpwood, round and split, coniferous | 316 | 0.6 | 189. | 1.852 | 551.7 | 30 | 30 | 30 |
| Wood fuel, coniferous | 238 | 0.6 | 143. | 1.852 | 415.7 | 30 | 2 | 16 |
| Other industrial roundwood, non-coniferous | 114 | 0.70 | 80.0 | 1.767 | 221.9 | 50 | 100 | 75 |
| Other industrial roundwood, coniferous | 42 | 0.6 | 24.9 | 1.852 | 72.5 | 30 | 100 | 65 |
|  | 3904.76 | 0.65 | 2603 |  | 7326.2 | 43.8 | 18.5 | 31.2 |

## References

1. Balesdent J. et Recous S., 1997. Les temps de résidence du carbone et le potentiel de stockage de carbone dans quelques sols cultivés français. Can. J. Soil. Sci., 77, pp.187193

2. Basher T. and Akter F., 2022. The Role of Plants in Carbon Sequestration: Mechanisms, Ecosystem Contributions, and Their Impact on Mitigating Climate Change, Australian Herbal Insight, 5(1),1-5, 9943. 10.25163/ahi.519943

3. Blume H. et al., 2015. Scheffer/schachtschabel soil science. Springer Berlin Heidelberg. Doi: 10.1007/978-3-642-30942-7

4. Chen C. et al., 2019. China and India lead in greening of the world through land-use management. Nat. Sustain. 2, 122–129 (2019).

5. Chen S. et al., 2013. A new estimate of global soil respiration from 1970 to 2008. Chin Sci Bull, doi: 10.1007/s11434-013-5912-1

6. FAO, 2025. Fishery and Aquaculture Statistics – Yearbook. Rome. 10.4060/cd6788en;

7. FAO, 2018. Rice Market Monitor. Volume XXI, Issue No. 1, April 2018. https://openknowledge.fao.org/server/api/core/bitstreams/07b6c7b4-065b-47ac-88d5-bbdafcb2baae/content

8. FAO (n.d.). FAOSTAT Data. https://www.fao.org/faostat/en/#data

9. Friedlingstein P. et al., 2024. Global Carbon Budget 2023. Earth Syst. Sci. Data, 15, 5301–5369. Doi: 10.5194/essd-15-5301-2023

10. Friedlingstein P. et al., 2025. Global Carbon Budget 2024. Earth Syst. Sci. Data, 15, 5301–5369. Doi 10.5194/essd-17-965-2025

11. Grassi G. et al., 2023. Harmonising the land-use flux estimates of global models and national inventories for 2000–2020, Earth Syst. Sci. Data, 15, 1093–1114, Doi 10.5194/essd-15-1093-2023

12. Gundersen P., 2021. Old-growth forest carbon sinks overestimated. Nature. Doi: 10.1038/s41586-021-03266-z

13. Hay R., 1995. Harvest index: a review of its use in plant breeding and crop physiology. Annals of Applied Biology, 126: 197–216.

14. ING, 2013. Les flux de bois en forêt. https://inventaire-forestier.ign.fr/IMG/pdf/flux2016.pdf, 2006a. Guidelines for National Greenhouse Gas Inventories. Chapter 2: Stationary combustion.

15. IPCC, 2006. Guidelines for National Greenhouse Gas Inventories. Chapter 5: Cropland.

16. IPCC, 2019. Refinement to the 2006 IPCC Guidelines for National Greenhouse Gas Inventories. Chapter 5: Cropland.

17. IPCC, 2021: Summary for Policymakers. In: Climate Change 2021: The Physical Science Basis. Contribution of Working Group I to the Sixth Assessment Report of the Intergovernmental Panel on Climate Change [Masson-Delmotte, V., P. Zhai, A. Pirani, S. L. Connors, C. Péan, S. Berger, N. Caud, Y. Chen, L. Goldfarb, M. I. Gomis, M.

18. Jia R. et al., 2025. A large global soil carbon sink informed by repeated soil samplings. bioRxiv preprint. Doi: 10.1101/2025.04.25.650716

19. Karadavut U. et al., 2008. A Growth Curve Application to Compare Plant Heights and Dry Weights of Some Wheat Varieties. American-Eurasian J. Agric. & Environ. Sci., 3 (6): 888–892

20. Keeling C. et al., 2001. Exchanges of atmospheric CO_2_ and ^13^CO_2_ with the terrestrial biosphere and oceans from 1978 to 2000. I. Global aspects, SIO Reference Series, No. 01-06, Scripps Institution of Oceanography, San Diego, 88 pages.

21. Koutsoyiannis D, Onof C, Kundzewicz ZW, Christofides A. 2023. On Hens, Eggs, Temperatures and CO_2_: Causal Links in Earth’s Atmosphere. Sci.; 5(3):35. 10.3390/sci5030035

22. Lal R., 2008. Sequestration of atmospheric CO_2_ in global carbon pools. Advance Article on the web. Doi: 10.1039/b809492f

23. Le Quéré C. et al., 2013. Global carbon budget 2013, Earth Syst. Sci. Data Discuss., 6(2), 689–760, Doi: 10.5194/essdd-6-689-2013.

24. Luyssaert S. et al., 2008. Old-Growth Forests as Global Carbon Sinks. 10.1038/nature07276

25. Luyssaert S. et al., 2021. Reply to: Old-Growth Forests Carbon Sinks Overestimated. Nature, 591, 24–25. 10.1038/s41586-021-03267-y

26. Ma S., He F., Tian D. et al., 2018. Variations and determinants of carbon content in plants: a global synthesis, Biogeosciences, 15, 693–702, 10.5194/bg-15-693-2018

27. Met Office Hadley Centre, 2022. 2021 continues warm global temperature series, Grahame Madge. https://www.metoffice.gov.uk/about-us/news-and-media/media-centre/weather-and-climate-news/2022/2021-hadcrut5-wmo-temperature-statement

28. Mignot A. et al., 2025. Sea Surface Temperature and Wind Extremes: Their Impact on Decadal Ocean Carbon Sink Variability. Preprint. Correspondence to

29. Mottet A. et al., 2017. Livestock: On our plates or eating at our table? A new analysis of the feed/food debate.Global Food Secur. 14, 1–8. Doi: 10.1016/j.gfs.2017.01.001

30. Muller-Feuga A., 2024a. The Recognition of Carbon Capture and Storage by Plants, Journal of Agricultural Science, 16, 7. Doi: 10.5539/jas.v16n7p1

31. Muller-Feuga A., 2024b. Plant Cultivation: A Strong and Sustainable Response to CO_2_ Emissions. Journal of Geoscience and Environment Protection, 12, 68–88. Doi: 10.4236/gep.2024.1210005

32. Muller-Feuga A., 2025a. The 50-Year CO_2_ Balance: Crucial Roles of Agriculture, Forestry and the Ocean. Great Britain Journals Press, Vol 24 Issue 15. Doi:10.34257/LJRSVOL24IS15PG1

33. Muller-Feuga A., 2025b. Reappraisal of the Place of Cultivated Plants in the World Carbon Budget. Voice of the Publisher, 11, 412–441. 10.4236/vp.2025.113029

34. Nissan A. et al., 2023. Global warming accelerates soil heterotrophic respiration. Nature Communications 14:3452. Doi 10.1038/s41467-023-38981-w

35. NOAA, n.d. The Surface Ocean CO_2_ Atlas. https://www.pmel.noaa.gov/co2/story/SOCAT

36. Pan Y. et al., 2024. The enduring world forest carbon sink. Nature 631, 563–569 (2024). Doi : 10.1038/s41586-024-07602-x

37. Pellerin S. et al., 2019. Stocker du carbone dans les sols français, Quel potentiel au regard de l’objectif 4 pour 1000 et à quel coûtSynthèse du rapport d’étude, INRA (France), 114 pp.

38. Raich J. W., Potter C. S. and Bhagawati D., 2002. Interannual variability in global soil respiration, 1980-94. GlobalChange Biology 8:800–812. doi: 10.3334/CDIAC/lue

39. Richet P., 2021. Le climat et la relation température-CO_2_. Un réexamen épistémologique du message des carottes glaciaires. Institut de Physique du Globe de Paris, 1 rue Jussieu, 75005 Paris, France. Traduction d’un article publié dans History of Geo- and Space Sciences, 12, 97–110 (2021)

40. Ritchie H., Rosado P., Roser M., 2023. CO₂ and Greenhouse Gas Emissions. Global Change Data Lab. https://ourworldindata.org/co2-and-greenhouse-gas-emissions

41. Saugier B., Roy J., Mooney H., 2001. Terrestrial global productivity. London: Academic Press.US$99.95 (hardback). 573 pp.

42. The Mainichi, 2025. A glimpse into how Japanese gov’t stores 910,000 tons of rice while preserving quality. Japanese original by Satoshi Fukutomi, Business News Department.

43. Tieszen L. et al., 1983. Fractionation and turnover of stable carbon isotopes in animal tissues: Implications for ∂^13^C analysis of diet. Oecologia, Springer-Verlag, 57:32–37.

44. Tijero V. et al., 2021. Fruit Development and Primary Metabolism in Apple. Agronomy 2021,11,1160. Doi: 10.3390/agronomy11061160

45. United Nations, 2024. Department of Economic and Social Affairs, Population Division.

46. World Population Prospects 2024: Methodology of the United Nations population estimates and projections. UN DESA/POP/2024/DC/NO. 10, July 2024 [Advance unedited version]

47. Vanclay JK., 1994. Modelling forest growth and yield : applications to mixed tropical forests, CAB International, Wallingford, UK.

48. Vidal E. et al., 2020. Sustainable forest management (SFM) of tropical moist forests: the case of the Brazilian Amazon. In Achieving sustainable management of tropical forests. Eds

49. Wang J. et al., 2017. Soil and vegetation carbon turnover times from tropical to boreal forests. Funct Ecol. 32: 71–82. Doi: 10.1111/1365-2435.12914

50. Wolf J. et al., 2015. Biogenic carbon fluxes from global agricultural production and consumption, Global Biogeochem. Cycles, 29, 1617–1639, Doi: 10.1002/2015GB005119.

